# Different methods to estimate the phase of neural rhythms agree, but only during times of low uncertainty

**DOI:** 10.1101/2023.01.05.522914

**Authors:** Anirudh Wodeyar, François A. Marshall, Catherine J. Chu, Uri T. Eden, Mark A. Kramer

**Author notes:** Correspondence should be addressed to Mark A. Kramer.

## Abstract

Rhythms are a common feature of brain activity. Across different types of rhythms, the phase has been proposed to have functional consequences, thus requiring its accurate specification from noisy data. Phase is conventionally specified using techniques that presume a frequency band-limited rhythm. However, in practice, observed brain rhythms are typically non-sinusoidal and amplitude modulated. How these features impact methods to estimate phase remains unclear. To address this, we consider three phase estimation methods, each with different underlying assumptions about the rhythm. We apply these methods to rhythms simulated with different generative mechanisms and demonstrate inconsistency in phase estimates across the different methods. We propose two improvements to the practice of phase estimation: (1) estimating confidence in the phase estimate, and (2) examining the consistency of phase estimates between two (or more) methods.

**Significant Statement:** Rhythms in the brain can coordinate the activity of individual neurons and communication within brain networks, making these rhythms a target for therapeutic interventions. Brain rhythms manifest in diverse ways, appearing with sinusoidal and non-sinusoidal waveforms, multiple peak frequencies, and variable durations. Across this diversity of rhythms, an important feature to characterize is the rhythm’s phase. In this manuscript, we demonstrate the ambiguity inherent in estimating phase from neural data. We propose estimating uncertainty in the phase and comparing multiple phase estimators improves phase estimation in practice.

## Introduction

A rhythm in the local field potential (LFP), magneto/electroencephalogram (M/EEG), or intracranial EEG (iEEG) is a waveform repeating without overlap. A cycle of a rhythm is an expression of the waveform (Cole & Voytek, 2017). Once the waveform is defined the phase can be used as a local time index of a cycle relative to the waveform. The phase of rhythms may have a role to play in perception (Busch et al., 2009; Herbst et al., 2022; Fiebelkorn et al., 2018; Helfrich et al., 2018; VanRullen, 2016), communication across neural populations (Fries, 2015; Roberts et al., 2013; Wodeyar et al., 2022), co-ordination within a neural population (Canolty & Knight, 2010) and co-ordination between local field potentials and neural spikes (Ray et al., 2008; Zanos et al., 2012; Jarvis & Mitra, 2001; Lepage et al., 2011; Pesaran et al., 2018). Phase has been of critical importance for therapeutic interventions as well (Widge & Miller, 2019): e.g., acoustic stimulation during sleep can induce slow waves (Ngo et al., 2013) and deep-brain stimulation can reduce Parkinsonian tremor (Cagnan et al., 2017).

In practice, phase is estimated with a Fourier analysis on tapered versions of the time series (Lepage et al., 2013), such as by using a Hilbert transform. In this approach, we assume that the time series consists of a stable rhythm accurately described as a harmonic (single frequency) or a narrowband waveform. A truly narrowband signal has spectral power concentrated in a tight frequency band. Any cycle will have an average rate of change of phase equal to the center frequency of the frequency band of maximum spectral power. This assumption determines the waveform and the phase. However, specifying phase for a cycle is difficult when the rhythm is inseparable from noise, the waveform has non-sinusoidal properties (Jones, 2016), or there is distortion of the waveform (such as from filtering a time series (Yael et al., 2018)).

Recent evidence suggests that a narrowband model for neural rhythms may be inaccurate. For example, visual cortex gamma band activity (30-90 Hz, c.f. Buzsaki 2006) is amplitude modulated and lacks narrowband periodicity (Burns et al., 2010; Spyropoulos et al., 2022; Xing et al., 2012). In this case, stochastically-driven damped harmonic oscillators may better model the gamma activity. The discretized version of such a model is an autoregressive process of order two (abbreviated as AR(2), see Appendix A of Kramer, 2022). Spyropoulos et al. (2022) show that an AR model fits the broadband power spectrum of the gamma rhythm well. AR models for both non-invasive (Fran & Bilo 1985) and invasive (Nozari 2020) electroencephalogram recordings support a broadband representation for brain rhythms. In addition, several studies suggest that brain rhythms can consist of non-sinusoidal waveforms (Bernardi et al., 2019, Cole & Voytek 2017, Cole & Voytek 2018, Ouedraogo 2016), exhibit multiple nearby peak frequencies (Tort 2010), and persist for short periods at a time (Jones 2016, van Ede 2018). These observations suggest brain rhythms may be broadband, and in these cases narrowband models lead to phase estimates which do not capture what is intuitively sought by the practitioner.

Phase is relatively defined, and how we estimate phase depends on the interpretation sought by the practitioner. For any simulated time series without an explicit phase parameter, phase must be independently estimated, thus there is no “true phase”. In this manuscript, we consider how different methods define phase when each is applied to simulated signals generated from different stochastic processes, each time series lacking an explicit phase parameter. Through this analysis, we identify moments when phase estimates agree and disagree across methods and examine how each method performs in different simulation scenarios. We consider three distinct phase estimators: (i) the Hilbert transform of a narrowband filtered time series, (ii) a state space model of rhythms, and (iii) the Poincaré section. We apply each of these methods to three simulated time series: (i) a broadband rhythm in pink noise, (ii) an AR(2) process, and (iii) a Fitzhugh-Nagumo oscillator. Comparing the cross-method consistency in phase, we show how uncertainty metrics can be used to improve consistency; when any method shows confidence in the phase estimate, the different methods tend to agree. Finally, we illustrate example applications of these methods to estimate coherence and phase-amplitude coupling in *in vivo* data. We show how estimating uncertainty in the phase helps clarify the link between phase and other quantities of interest. Understanding the role of phase in neural dynamics requires us to be aware that phase is not unique and phase estimates may vary depending upon the method employed.

## Methods

### Overview of Phase Estimation Approaches

We apply three methods to estimate phase, each of which makes different assumptions for the rhythm model. The first method we consider consists of bandpass filtering the data and then estimating the phase via the Hilbert transform. While a common method in neuroscience applications (e.g., Canolty & Knight, 2010; Navas-Olive et al., 2020; Petersen & Buzsáki, 2020; Zhong et al., 2017), this approach is sensitive to noise and susceptible to waveform distortion. The second method we consider is a state space model with the state, representing the amplitude and phase, modeled by stochastically-driven oscillators. The state space model makes stronger assumptions of the data but is more resistant to model noise when the model class optimally explains the periodicity in the data. The final method we consider is the application of the Poincaré section, which is the most general method considered but is also the most sensitive to noise and user-defined choices. A complication is that different models which fit a brain rhythm equivalently well can still give phase estimates tracking distinct features of a rhythm. In what follows, we use this ambiguity when specifying phase to explore approaches to validate analyses involving phase.

### Finite Impulse Response filter and Hilbert Transform (FIR-Hilbert) Method

Fourier-transform based analysis, a wavelet-transform, or a Hilbert transform employ essentially the same procedure (Bruns, 2004) to estimate the phase but with different tapering functions. Critically, all these approaches demarcate a set of frequencies (bandwidth of the narrowband) as relevant to capture the neural signal of interest and a set of frequencies as being irrelevant (for example, filtering to select the alpha band, defined as 8 – 13 Hz). This demarcation is intended to separate the signal and noise without relying on a parameter to explicitly model the noise contribution (Donoghue et al. 2020). Given the essential equivalence of the windowed-Fourier, wavelet, and Hilbert transforms (Bruns, 2004), we focus here on the Hilbert transform; i.e., we apply a filter to the data time series to select a frequency band of interest, apply the Hilbert transform, and derive the instantaneous phase (Chavez et al., 2006).

For an interpretable instantaneous phase - where the phase captures the waveform periodicity as opposed to amplitude modulation - the Hilbert transform (or a windowed-Fourier transform or a wavelet-transform) should only be applied when the spectral support (i.e., band for which spectral mass is non-zero) of the amplitude time series does not overlap with the spectral support of the phase time series (Boashash, 1991; Chavez et al., 2006). This is a consequence of the Bedrosian theorem (Bedrosian, 1963; see Boashash, 1991 for a discussion) which shows:

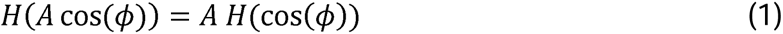

Where *H* is the Hilbert transform operator, *A* is the amplitude time series, and *ϕ* is the phase time series. The Bedrosian theorem holds when the effective spectral support of *A*, i.e., the frequencies with maximal spectral mass, are distinct from the effective spectral support of *ϕ*. Thus, any data time series with multiple rhythms or a rhythm with amplitude modulations may lead to phase estimates under the Hilbert transform that are difficult to interpret as the argument of a cosine function; see Chavez et al., 2006 for extended discussions of these theorems.

Another factor affecting the interpretability of the phase under the Hilbert transform is the parameter set used for the narrowband filter. A practitioner must decide the filter type, filter order, and window duration (see Widmann et al. (2015) for a discussion and Zrenner et al. (2020) for an analysis of different parameter choices). The practitioner must also choose the filter’s bandwidth; any waveform whose spectral support is wider than the bandwidth of the filter will be altered in the filtered time series.

For all analyses here, we apply a least-squares linear-phase FIR filter of order 3 times the longest cycle length expected (as in, e.g., Cohen, 2008 and Nadalin et al., 2019). After forward and backward filtering of the data (using the filtfilt function in MATLAB), we apply the Hilbert transform to the data to estimate the quadrature component of the signal. We compute the analytic signal using the hilbert function in MATLAB, which computes the quadrature component (the imaginary term) and combines it with the in-phase component (the real term). Finally, we estimate phase using the four-quadrant arctan function implemented by the angle function in MATLAB.

We estimate the 99% confidence interval for the FIR-Hilbert phase estimator, assuming stationary noise and complex-valued normality for the additive noise to the analytic signal, using (Eq. 3.18 of Lepage et al., 2013):

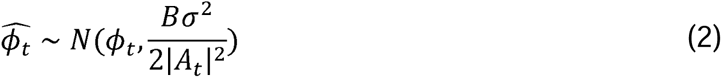

Where *A_t_* is the time-*t* element of the amplitude time series, *B* is the bandwidth of the filter divided by sampling frequency, and *σ*^2^ is the additive noise variance, which is estimated using the residual after removing the filtered data from the unfiltered data. This confidence interval is derived by operationalizing the filter and Hilbert transform in terms of linear operators applied to the signal, and assuming for the analytic signal a simple signal plus additive noise model (see Eqs 3.2 - 3.11 of Lepage et al., 2013), where the noise is complex-Gaussian. From the inferred confidence bounds, asymptotic expressions for the maximum likelihood estimate of phase can be derived, assuming that the amplitude of the signal is significantly higher than the standard deviation of the noise (for further discussion, see Lepage et al., 2013).

We note that the confidence interval for the phase estimate depends on the filter bandwidth (*B*): the wider the bandwidth, the greater the variance (or lower the confidence) in the phase estimate (Eq 2). Similarly, a filter with center frequency misaligned with the center frequency of the true signal will increase variance in the phase estimate by increasing the additive noise variance (*σ*^2^), the inferred variance of the data after removing the filter-applied signal) and reducing the estimated amplitude time series (*A_t_*, the amplitude time-series of the filter-applied signal). In this case, increasing the filter bandwidth (*B*) may reduce variance in the phase estimate by reducing the additive noise variance (*σ*^2^) and increasing the estimated amplitude time series (*A_t_*). Thus, characterizing the tradeoff between filter bandwidth (*B*), and the additive noise variance (*σ*^2^) and estimated amplitude time series (*A_t_*) depends on the specific signal of interest.

### State Space Model

A state space modeling approach to specify phase (Matsuda & Komaki, 2017; Wodeyar et al., 2021; Soulat et al., 2022) expects the random time series to exhibit periodicity similar to that of a vector-valued AR(2) process. We assume that the analytic signal of a rhythm is a latent, unobservable process, that we can estimate from the observed data. We build this assumption into a state space structure, with the state equation (see Eq. 3 below) representing the latent analytic signal and the observation equation (see Eq. 5 below) representing the observed data. On receipt of each new sample of the observed data, we correct the estimate of the state using a Kalman filter - an optimal strategy when the state update equation is linear and Gaussian (see Chapter 2 of Schiff, 2012).

To illustrate this approach, we consider the case of tracking a single rhythm whose central baseband frequency is 6 Hz. The state equation of the univariate analytic signal *x* for this case is:

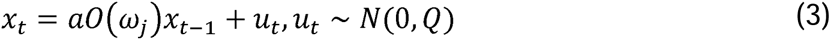

where *x_t_* is a real-valued 2-D signal with real (Re(*x_t_*)) and imaginary (Imag(*x_t_*)) parts. The state equation yields a prediction for the current state (*x_t_*) by rotating realized values of the state (*x_t_*_-i_) in the complex plane and adding the Gaussian innovation *u_t_*. The rotation matrix O(ω) is given by:

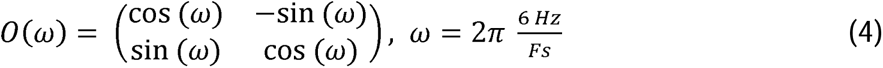

where *F_s_* is the sampling frequency in Hertz. After rotation of each realized value of *x_t_*_-1_, the resulting output is damped by a factor *a ∈* [0,1), and then a realization of the Gaussian innovation (*u_t_*, with variance Q) is added. We estimate the observable at time *t* (*y_t_*) as the real part of the state *x_t_* at time *t* in the presence of additive observation noise:

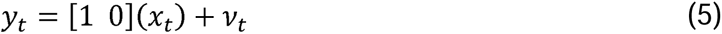

where *v_t_* is the *t^th^* element of Gaussian IID noise (*ν_t_* ∼ *N* (0, *σ*^2^)).

When applying this method to identify brain rhythms and track the associated analytic signal, we need to estimate *Q*, *a*, *σ*^2^, *ω* (assumed 6 Hz in illustration above). Estimating these parameters can be done by applying the expectation-maximization algorithm (Shumway & Stoffer, 1982), and depends on good initial estimates for the parameters (see Matsuda & Komaki, 2017 and Soulat et al., 2022 for potential initialization strategies). We use the model described in Equations (3) - (5) under a Kalman filter framework, after estimating parameters, to obtain the optimal state estimates for each observed data sample. Further, we perform a backward smoothing operation (Shumway & Stoffer, 1982; Soulat et al., 2022), i.e., sending a time-reversed version of the data through the filter. Finally, after smoothing, for each data sample we derive the posterior distribution for the state prediction that we use to define the credible interval over the phase (Morey et al., 2016). We sample 10,000 times from the posterior distribution for the state and use this to extract 99% credible intervals (see *Methods* in Wodeyar et al., 2021 for details).

### Poincaré Section Method

For any K-dimensional dynamical system and associated particle trajectory, we can define a Poincaré section as a K-1 dimensional plane which intersects the considered trajectory. We specify phase relative to the moment of plane crossing. For example, in a one-dimensional system (i.e., the activity varies along a single axis), negative to positive zero-crossings define a Poincaré section, where the origin is the ordinate through which we mark the particle trajectory. As described by Chavez et al., 2006 and others (Kralemann et al., 2007; Rosenblum et al., 1997), phase specified using a Poincaré section depends upon the selection of observables (for example, the signal alone versus a time-delayed embedding (Takens, 1981)), and how the section is defined. In this manuscript, we consider K=2 and select zero-crossings to define the phase. From one zero-crossing (i.e., from the voltage traveling from negative to positive values) to the next, 2π radians are uniformly covered by the evolution of the phase. We denote *t_n_* and *t_n_*_+1_ as the times of the *n*^th^ and *n*+1^th^ zero-crossings of the time-evolving particle. During [*t_n_*, *t_n_*_+1_) the phase-space of the particle completes one full rotation. The phase *ϕ* is specified by a linear ramp on [*t_n_*, *t_n_*_+1_) (Rosenblum et al., 1997):

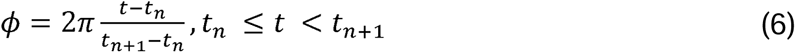

For a pair of consecutive zero-crossing times, the Poincaré section assigns a phase of 0 at *t_n_* and 2Π at the sample immediately prior to *t_n_*_+1_. We note that, in the other two methods of phase specification, 0 and 2Π are assigned to phase at the amplitude maxima. To make the phase mapping isomorphic across methods, we unwrap the phase (unwrap function in MATLAB) assigned by the Poincaré section, add Π/2, and wrap (wrapToPi in MATLAB) the phase to [-Π,Π]. This transformation shifts the phase by Π/2 and remaps phase to [-Π,Π]. Unlike the other two methods considered here, the Poincaré section does not explicitly model a source of noise and thus lacks a measure of uncertainty in the phase estimate. Despite this and as we will see in the *Results*, for non-sinusoidal rhythms, the Poincaré section serves as a useful comparison.

### Signal-to-Noise Ratio

We define the signal-to-noise ratio (SNR) as the extent to which the power within a limited band of frequencies around the central frequency is greater than power at other frequencies. We control SNR in our simulations to exceed 2.5 for clear visual identifiability of a rhythm. To infer the SNR for an oscillation in the data with center frequency in *F*, we apply:

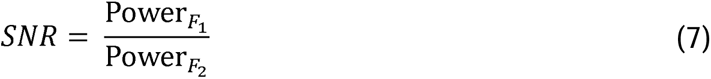

where ***F_1_*** is the frequency band of interest and ***F_2_*** are frequencies outside the frequency band of interest. Power*_F1_* is the spectral power across the frequency band ***F_1_*** as derived from a multitaper spectral estimator with 9 tapers using the mtspectrumc function from Chronux <u>(Bokil et al., 2010)</u>. In this manuscript, the summation in the numerator includes frequencies from 1 Hz below to 1 Hz above the central frequency, and the summation in the denominator includes frequencies 2 Hz below and 2 Hz above the central frequency in the denominator.

### Estimating Phase Difference Variability

To determine the consistency of phase across methods, we compute the circular standard deviation (Zrenner et al., 2020). This method characterizes the width of the distribution of phase differences between a pair of phase estimators around a mean difference in the phase. To measure the circular standard deviation of the phase differences between phases *θ_M_* and *θ_n_* for estimators *M* and *N*, we apply (Zrenner et al., 2020),

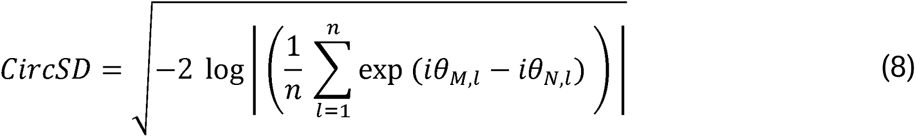

We note that this method of assessing phase across methods presumes that any bias (i.e., a phase difference centered at a value other than zero) is consistent across time (Zrenner et al., 2020). We interpret the *CircSD* as a measure in radians. We note that this interpretation is valid only when the spread of phase-difference angles is non-uniform; closer to a uniform distribution, this measure exceeds *2π*. However, for the cases considered here, this situation does not occur.

### Analysis of in vivo data

To illustrate application of the phase estimation methods, we consider an open-source dataset consisting of recordings from two invasive EEG electrodes (iEEG, 400 s sampled at 1000 Hz) in a non-human primate anesthetized with propofol (available at http://neurotycho.org/expdatalist/listview?task=75). Propofol anesthesia yields a brain state with synchronized brain rhythms (Lewis et al., 2012), resulting in strong cross-cortical coherence and phase-amplitude coupling relationships, phenomena proposed to coordinate neural spiking within and between brain regions (Fries, 2015; Jarvis & Mitra, 2001; Lepage et al., 2011; Pesaran et al., 2018). Here we apply different phase estimation methods to analyze the coherence in the slow delta band [0.25, 1.5 Hz] between the two recordings, and phase-amplitude coupling between the slow delta and beta [13, 30] Hz (defined from the classical range provided in (Engel & Fries, 2010)) bands within each recording (Ma et al., 2019).

To estimate the phase of slow delta activity via the FIR-Hilbert method, we apply an FIR filter to isolate the frequency band of interest (slow delta) and calculate the Hilbert transform to estimate the slow delta phase. We apply the same FIR-Hilbert approach to estimate the amplitude time series for the beta band (Cohen, 2008; Nadalin et al., 2019). For the state space method, we first estimate model parameters as well as the central frequencies for each band (slow delta and beta) using a subset of the data (10 s). We do so for each recording separately, and then use these parameters for state estimation across all 400 s of data (see Equations 6 - 10 in Wodeyar et al., 2021). For the Poincaré method, we first broadband filter the iEEG between 0.1 Hz to 50 Hz using a 4th order Butterworth filter. Filtering before applying the Poincaré method reduces the influence of higher frequency noise and slow drifts on zero-crossings. The Poincaré method does not yield an amplitude time series, so we assume the amplitude is constant and equal to 1. We note that, in this case, coherence values using the Poincaré method are akin to estimating a phase locking value (Lachaux et al., 1999).

We use phase estimates from each method to estimate the coherence of the slow delta [0.25 - 1.5 Hz] rhythm between the two recordings. Coherence is an estimate of the consistency in the phase and amplitude at one frequency (or in a single frequency band) between two electrophysiological time series (for a discussion see Pesaran et al., 2018). We compute coherence (*C*) for two complex-valued row vectors over time *E*_1_ and *E*_2_, each representing the amplitude and phase in one frequency band using the function corr in MATLAB.

Phase-amplitude coupling represents how the phase of a low frequency rhythm (here, the slow delta band) coordinates the amplitude of a high frequency rhythm (here, the beta band) (Hyafil et al., 2015). We note that we cannot apply the Poincaré method for phase-amplitude coupling estimation as this method does not estimate an amplitude time series. We therefore analyze the phase-amplitude coupling within each recording using only the Hilbert and state space methods. To visualize the phase-amplitude coupling, we divide the phase between 0 to 2*π* into 20 non-overlapping intervals and plot the average amplitude for each interval of phase (Canolty & Knight, 2010; Tort et al., 2010).

### Code Accessibility

All code developed for and applied in this paper are available for reuse and further development at: https://github.com/Eden-Kramer-Lab/UncertaintyInPhase. All code was run on a Dell XPS 15 7590 running Windows 10 and having 32 GB of RAM. A README file is included with the code that has a legend for all files.

## Results

A pure sinusoid is the only function that has an explicitly specified phase parameter, but it is too restrictive a model for most brain rhythms (Cole & Voytek, 2017). For the simulated time series considered below, an optimal specification of phase does not exist. The generative models used to simulate the time series have no explicit phase parameter, thus, phase estimates must be derived as functions of the generative stochastic processes. This precludes a “true phase”-i.e., a single phase parameter - and error metric. Because in practice only observations of a pure sinusoid have a true phase, we instead consider different methods to estimate phase, each imposing different assumptions about the data. We then compare the relative performance of these different methods. In what follows, we show that when any method indicates reduced uncertainty in the phase estimate, then all three methods tend to agree allowing practitioners to limit the phase estimates required to be considered.

We analyze simulation data from four generative models. First, we examine a case where the generative model has a phase parameter but is unlikely to be a good model of empirical data. Then, we consider three more empirically valid models: a non-parametric model, a simple stochastic parametric model, and a dynamical systems model. The first generative model can fit the broadest range of neural data, while the second and third generative models are relevant for smaller subsets of data. Each generative model serves as a potential exemplar of a range of empirical data. For all generative models, we make three assumptions. First, we assume that a rhythm always exists in the data with high signal-to-noise ratio (SNR, with SNR >= 2.5 to allow for an identifiable rhythm, see *Methods*). Second, we assume that the data consists of a single rhythm of interest. Third, we assume that any temporal variability in the data is derived from the generative model.

### Amplitude Modulated Sinusoid in Pink Noise

To consider the scenario of a well-defined, ground truth phase, we simulate a 6 Hz sinusoid with amplitude modulation at a rate of 0.56 Hz, such that every 1.8 s the 6 Hz sinusoid emerges for 0.9 s (see spectrum to understand the signal-to-noise level in Figure 1a and example trace in Figure 1b). In this signal, a meaningful phase parameter exists only during times when the sinusoid is present. To this signal, we add pink noise (power proportional to 1/f^1.5^; Figure 1a) with random starting phases for the sinusoid at each frequency bin. We analyze 1000 instantiations of these simulated data, each of duration 10 s. This signal represents the burst-like dynamics of rhythms in brain electrophysiological time series.

**Figure 1:**
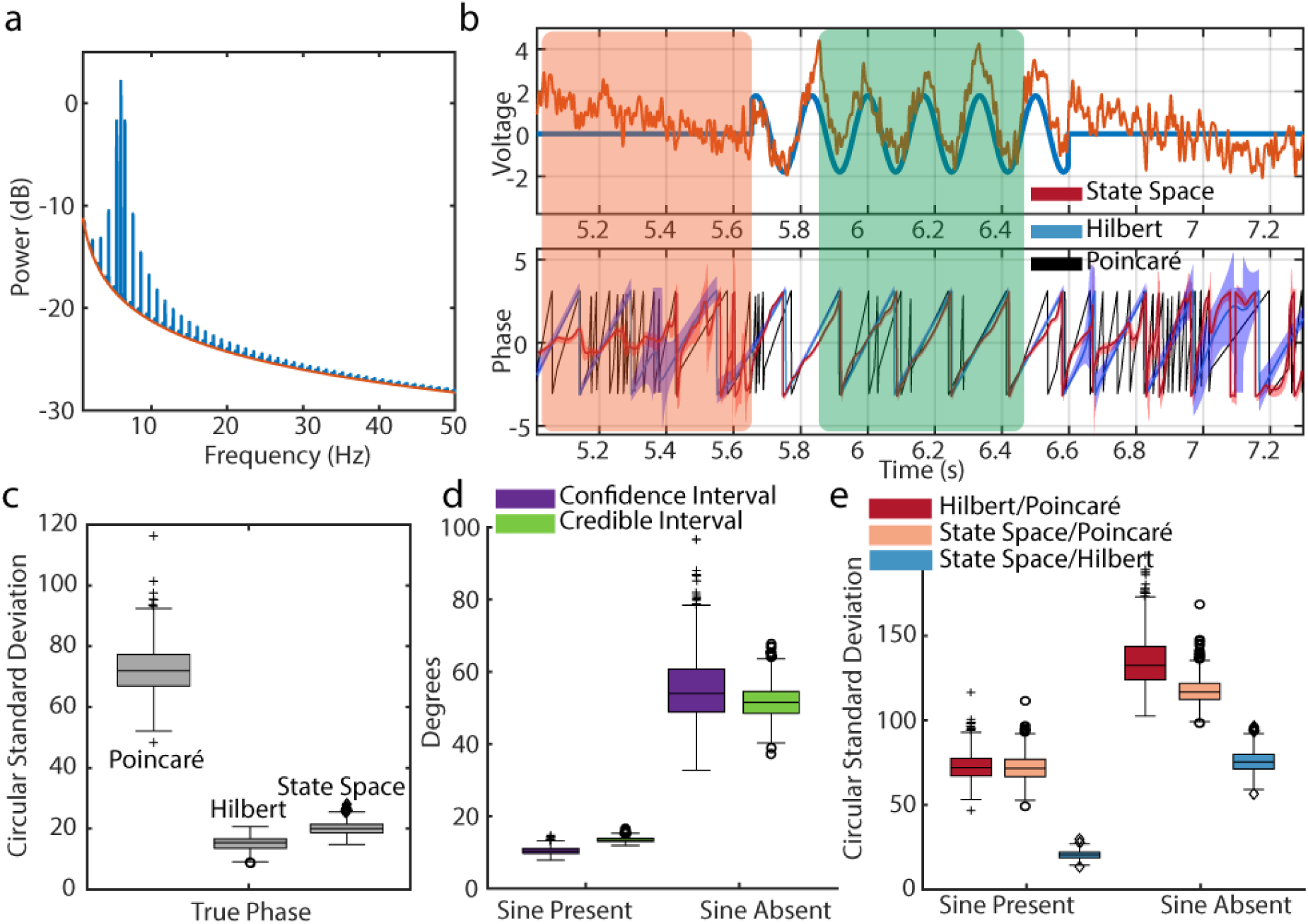
FIR-Hilbert and state space methods more accurately recover true phase for an amplitude modulated sinusoid in pink noise. (a) Power spectrum of the background pink noise (red) and spectral peak at 6 Hz (blue) as well as harmonics from square-wave amplitude modulation. (b) (Top) Example trace of the 6 Hz amplitude modulated sinusoid (blue) and the summated data with pink noise (orange). Rectangles indicate periods with (green) and without (red) the sinusoid. (Bottom) Phase estimates from the Poincaré method (black), FIR-Hilbert method (mean estimate blue, confidence intervals shaded blue) and state space method (mean estimate red, credible intervals shaded red). (c) Distributions of differences in the estimated phase across 1000 simulations between each method and the true phase during times when the sinusoid is present. (d) Confidence and credible interval distributions across 1000 simulations when the sinusoid is present and absent. (e) Distributions of differences across 1000 simulations between methods when using time segments with or without the sinusoid. Estimates from all methods better align during segments of data when the sinusoid is present.

We estimate the phase of the simulated 6 Hz rhythm using the Poincaré method, the FIR-Hilbert method, and the state space method (examples in Figure 1b). Visual inspection suggests that the latter two methods produce similar phase estimates when the sinusoid is present. However, during periods when the sinusoid is not present (red rectangular box in Figure 1b), the FIR-Hilbert method computes phase estimates whose trajectory completes full cycles (continuous traversal from -*π* to *π*), while phase estimates under the state space method tend to concentrate near 0 or *π*. Visual inspection shows that the phase from the Poincaré method is affected by the sudden zero crossings, which arise from the appreciable spectral power at frequencies above 6 Hz. This leads to rapid evolution of the phase, suggesting cycles that may be undesirable when the goal is to track the phase of the 6 Hz sinusoid. These qualitative observations are confirmed by direct comparison of the phase estimates to the true phase when the 6 Hz sinusoid is present (Figure 1c); the state space and FIR-Hilbert methods more accurately (median circular standard deviation of 20 degrees and 15 degrees, respectively) track the true phase compared to the Poincaré method (median circular standard deviation of 72 degrees).

The state space and FIR-Hilbert methods both support the calculation of metrics of uncertainty in the phase estimate - credible intervals for the state space method and confidence intervals for the FIR-Hilbert method (see Methods). To compare these metrics of uncertainty, we calculate the distributions of the widths of the credible and confidence intervals for time points when the sinusoid is, and is not, present (Figure 1d). We find that both measures of uncertainty unambiguously distinguish when a sinusoid is present (the uncertainty is low, FIR-Hilbert median confidence interval width of 10 degrees; and state space method median credible interval width of 13 degrees) versus when a sinusoid is absent (the uncertainty is high, FIR-Hilbert median confidence interval width of 54 degrees and state space method median credible interval width of 51 degrees).

Finally, as we lack knowledge of the true phase in real data and in more realistic simulations of neural activity, we compare the phase estimates between methods as a proxy to judge when multiple characteristics of the signal used to define phase are consistent. Comparisons between all three methods and their metrics of uncertainty reveal when meaningful estimates of phase can be calculated, i.e., when the sinusoid is present. For each pair of methods, we compute the difference between phases and construct the circular standard deviation histogram of the phase estimate differences across time points when the sinusoid is present and when it is absent. We find that when the sinusoid is absent there is a greater cross-method difference in phase estimates, consistent with greater discrepancy in how the phase is defined. We conclude that for an amplitude modulated sinusoid the state space and FIR-Hilbert methods more accurately recover the true phase compared to the Poincaré method.

### Broadband Rhythm in Pink Noise

To match empirical observations in neuroscience (Miller et al., 2009a; Miller et al., 2009b), we consider a phenomenological model of rhythms with increased power over a broad range of frequencies, emerging above 1/f^α^ background activity (i.e., the background activity power decreases with frequency as 1/f^α^, where *α ∈* [1,3] (Donoghue et al., 2020). We simulate this type of data in the frequency domain as a complex-valued vector. To do so we add a Gaussian-function shaped broadband amplitude (center frequency 6 Hz, standard deviation 1 Hz) with a zero phase to a background of pink noise (power proportional to 1/f^1.5^; Figure 2a) with random phases at each frequency. The inverse discrete Fourier transform applied to this complex-valued vector yields a time series of 10 s duration.

**Figure 2:**
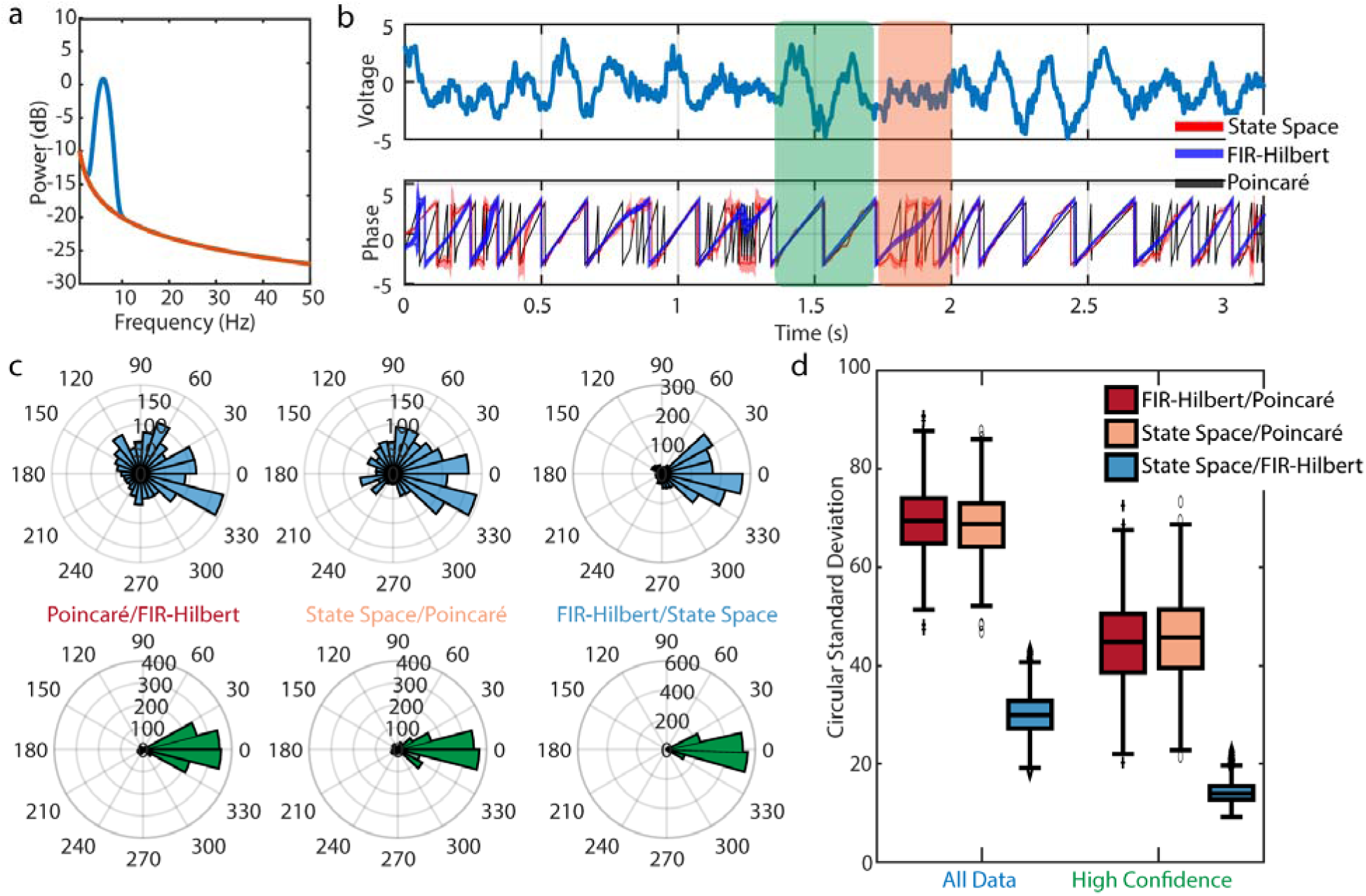
The FIR-Hilbert and state space methods produce consistent results for a broadband signal in pink noise. **(a)** Analytic power spectrum of the background pink noise (red) and broadband spectral peak at 6 Hz (blue). **(b)** (Top) Example trace of broadband signal with added pink noise. (Bottom) Phase estimates from the Poincaré method (black), FIR-Hilbert method (mean estimate blue, confidence intervals shaded blue) and state space method (mean estimate red, credible intervals shaded red). All three methods show periods where they disagree (t *∈* 1.75 s, see orange rectangle) and agree (t ∼ 1.5 s, green rectangle). **(c)** Distributions of differences in the estimated phase between each pair of methods using (upper) all example data or (lower) a subset of data with confident phase estimates. Estimates from all methods better align during segments of data with confident phase estimates. **(d)** Distributions of differences between methods across 1000 simulations. Using samples of data with high confidence in the phase improves consistency between methods. For broadband rhythms, the state space and FIR-Hilbert methods are more similar than the Poincaré method.

We estimate the phase of the simulated 6 Hz rhythm using the Poincaré method, the FIR-Hilbert method, and the state space method (examples in Figure 2b). Visual inspection suggests that the latter two methods produce similar phase estimates when the rhythm is visible; at these times the credible intervals of the state space method tend to overlap the confidence bounds of the FIR-Hilbert method (i.e., the two shaded curves tend to overlap in Figure 2b, see t ≈ 1.5 s). During periods of low amplitude (*t ∈* [1.75,2] *s*), the FIR-Hilbert method computes phase estimates whose trajectory completes full cycles (continuous traversal from -*π* to *π*) while phase estimates under the state space method remain consistently near ±*π*. Visual inspection of the phase estimated by the Poincaré method reveals a different result than what is seen using the other methods; sudden zero crossings, which arise because of appreciable spectral power at high frequencies, lead to rapid phase-series cycles that may be undesirable if the goal is to track phase for fluctuations at lower frequencies. The distributions of phase differences between all pairs of methods support the similarity between the state space and FIR-Hilbert methods and their contrast with the Poincaré method (Figure 2c); phase differences between the state space and FIR-Hilbert methods concentrate near 0, while phase differences with the Poincaré method appear at all angles. Repeating this analysis for 1000 realizations of the generating model and considering the variability of phase differences using circular standard deviation (see *Methods: Estimating Phase Difference Variability*), we find consistent results (Figure 2d, left boxplots).

We infer moments of consistent phase using the confidence bounds determined from the FIR-Hilbert method and the credible intervals from the state space method. We designate time segments with both smaller confidence bounds and smaller credible intervals as “high confidence” time segments. These are times when the values are jointly in the bottom quartile (as empirically determined) for both uncertainty metrics. Across high confidence time segments, the phase difference between the FIR-Hilbert and state space methods shows a circular standard deviation of 14.22 ± 0.07 degrees (mean ± standard error of the mean or SEM) reduced from 30.02 ± 0.13 degrees when considering all data (i.e., times with and without high confidence in the phase estimates). Similarly, the circular standard deviation between the FIR-Hilbert/Poincaré (44.8 ± 0.3 degrees) and state space/Poincaré (45.8 ± 0.3 degrees) methods decreases in high confidence time segments, compared to all data (FIR-Hilbert/Poincaré: 69.5 ± 0.2 degrees; state space/Poincaré: 68.6 ± 0.2 degrees; Figure 2d, right boxplots). However, we note that circular standard deviation remains high when comparing the Poincaré method for both choices of time segments (Figure 2d, red and pink boxplots).

We note that different filtering approaches may impact the FIR-Hilbert results. To investigate this, we also applied to these simulated data a variety of common filtering approaches (as in Zrenner et. al. 2020): windowed sinc and least squares finite impulse response filters of 2nd to 5th orders; Butterworth filters of order 2, 4, and 6; Chebyshev Type I filters of order 2, 4, and 6; and elliptical filters with 10 and 20 dB attenuation. We maintained a consistent passband across approaches to allow comparability and applied the Hilbert transform to estimate the phase. For each filtering approach, we then computed the phase difference with the primary FIR-Hilbert approach defined in Methods. Different filtering approaches produce different phase estimates, as expected (Figure 2e). However, the circular standard deviations between estimates from different filtering approaches are much smaller (median is less than 20 degrees in 15 out of 16 cases) than between the FIR-Hilbert Method and the state space or Poincaré method. Because these filtering approaches capture the narrowband peak in the simulated data, we find that different filtering methods produce similar phase estimates, consistent with Sameni & Seraj, 2017 and Zrenner et al., (2020). Different filtering approaches with the same central frequency and bandwidth nonetheless produce different phase estimates, as expected (Figure 3a). However, the circular standard deviations between estimates from different filtering approaches are much smaller (median is less than 20 degrees in 15 out of 16 cases) than the deviation in phase estimates from the FIR-Hilbert method and either of the state space or Poincaré methods (Figure 3b.i). Because the filtering approaches focus on capturing a narrowband peak in the simulated data, we find that different filtering methods produce similar phase estimates, consistent with Sameni & Seraj (2017) and Zrenner et al., (2020). Finally, we examine the impact of the center filter frequency on the FIR-Hilbert method. To do so, we adjust the center filter frequency between 3 Hz and 9 Hz, while keeping the bandwidth (4 Hz) and filter approach (finite impulse response) fixed. We find that, as the central frequency of the filter deviates from the correct central frequency, phase differences compared to the primary FIR-Hilbert approach rapidly increase (e.g., the phase difference between the 3 Hz filter and 6 Hz filter exceeds that between the FIR-Hilbert and Poincaré methods; Figure 3b.i, 3b.ii). This difference is reduced (by approximately 20 degrees) when examining the subset of data with confident phase estimates (i.e., when the FIR-Hilbert confidence intervals are small; Figure 3c.ii). We conclude that during periods of high confidence in the phase estimate (e.g., when the rhythm has higher amplitude, reducing the confidence interval width), the phase is consistently estimated (similar phase estimates across filters) even when the center filter frequency deviates from the true central frequency.

**Figure 3:**
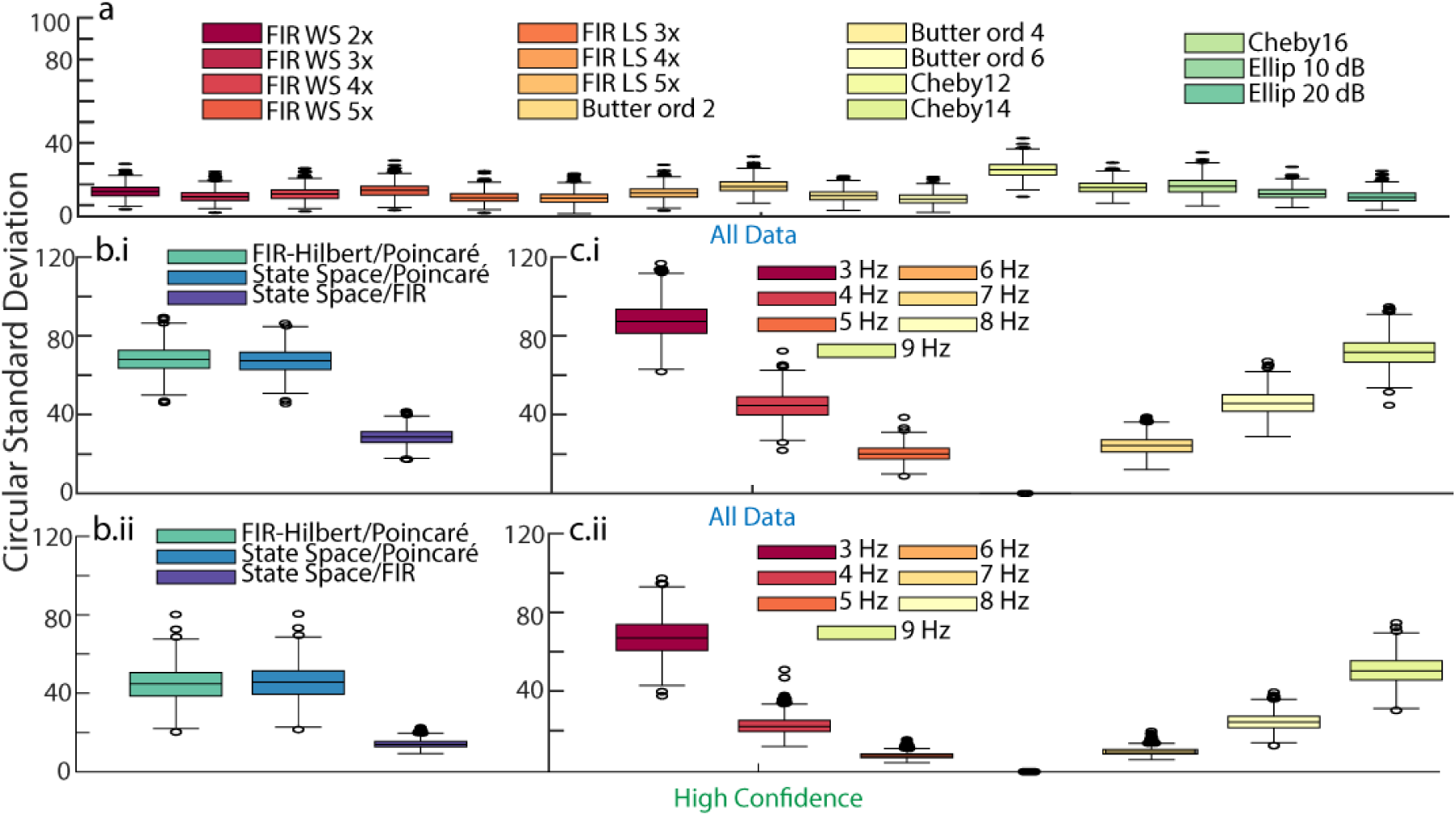
Different filter parameters minimally change FIR-Hilbert derived phase estimates. **(a)** Distributions of differences between Hilbert transform phase estimates with different filtering approaches (see legend) across 1000 simulations. The variability due to filtering approaches is less than the variability between methods. **(b)** Distributions of differences between phase estimation methods (i) for all data and (ii) when there is high confidence in the phase estimate. **(c)**Distributions of differences between Hilbert transform phase estimates with different central frequencies (+/− 2 Hz bandwidth, see legend) for (i) all data, and (ii) only data with high confidence in the phase estimate. Variability of phase estimates at different central frequencies rapidly increases away from the true central frequency (6 Hz).

We conclude that, for a broadband rhythm in pink noise, the state space and FIR-Hilbert methods produce comparable results. High frequency fluctuations in the time series produce intervals of rapid zero-crossings in the Poincaré method. Intervals exist with increased agreement between all three methods; these intervals occur when the state space and FIR-Hilbert methods produce confident estimates (i.e., small confidence bounds/credible intervals).

### Noise-driven Damped Harmonic Oscillator

Auto-regressive (AR) processes of order 2 are linear time-series models capable of exhibiting periodicity (Shumway & Stoffer, 2017). AR(2) processes sustaining a rhythm demonstrate spontaneous variability in the amplitude of the rhythm or amplitude modulation (Spyropoulos et al., 2022) and a 1/f^α^ proportional power spectrum. These features are consistent with features that characterize gamma band rhythms, as inferred from *in vivo* observations(Burns et al., 2010; Spyropoulos et al., 2022; Xing et al., 2012). Unlike the previous simulation where the additive noise is characterized by a 1/f^α^ power spectrum, here the Gaussian noise is passed through an AR(2) filter to yield spectra that has both a broadband peak centered at 6 Hz and a 1/f^α^ power spectrum; see analytic power spectrum in Figure 4a and an example trace in Figure 4b.

**Figure 4:**
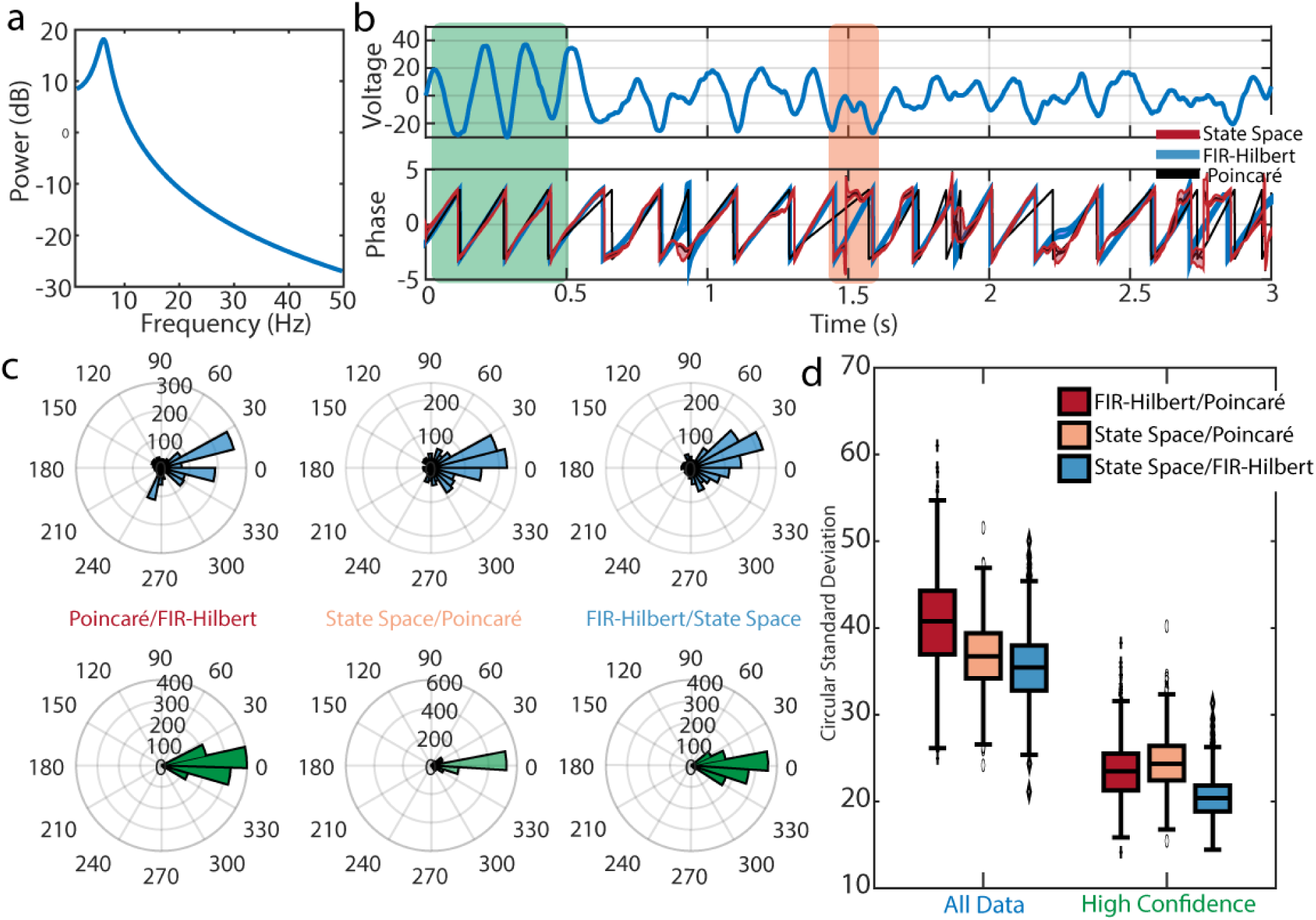
All three methods produce similar phase estimates for data generated by an AR(2) model. **(a)** Analytic power spectrum of the AR(2) model. A broadband spectral peak occurs at 6 Hz. **(b)** (Top) Trace of example AR(2) signal. (Bottom) Phase estimates from the Poincaré method (black), FIR-Hilbert method (mean estimate blue, confidence intervals shaded blue) and state space method (mean estimate red, credible intervals shaded red). Intervals occur in which the methods agree (t < 0.5 s) and disagree (t near 1.5 s, orange rectangle) **(c)** Distributions of differences in the estimated phase between each pair of methods using (upper) all example data or (lower) a subset of data with confident phase estimates. Estimates from all methods better align during segments of data with confident phase estimates. **(d)** Distributions of differences between methods across 1000 simulations. Using samples of data with high confidence in the phase improves consistency between methods.

Visual inspection of the estimated AR(2) phase reveals intervals of agreement and disagreement between the three methods (Figure 4b) as a result of amplitude modulation. For example, when the rhythm has high amplitude (*t* < 0.5 s), the uncertainty reduces and the three phase estimates are qualitatively similar in the features (e.g., peaks, troughs, and zero-crossings) tracked by the phase. At other times the methods disagree about what constitutes a full phase cycle. For example, at *t* near 1.5 s, the observed time series exhibits two low amplitude troughs. Between these troughs, the state space model suggests no phase cycle occurs (staying near ±*π*), the Poincaré method indicates a long phase cycle spanning beyond the two troughs, and the FIR-Hilbert method indicates a complete phase cycle between the two troughs. We find similar phase estimates across methods (Figure 4c, top row), with a circular standard deviation of 37.9 ± 0.1 degrees (mean and SEM across comparisons of all three methods over 1000 realizations of the simulation; see Figure 2d). Restricting analysis to high confidence time segments, we find increased consistency in phase estimates across methods (Figure 4c, bottom row) and a nearly 2-fold reduction in the circular standard deviations between methods (22.9 ± 0.1 degrees, 1000 realizations of the simulation; see Figure 4d).

We conclude that, for an AR(2) simulated time series, all three methods produce comparable results. Compared to the simulation of a broadband rhythm in pink noise (Figure 2), in the AR(2) simulated time series fewer high frequency fluctuations distort the phase estimates under the Poincaré method. Consistent with the previous simulation, when the state space method and FIR-Hilbert methods both produce confident estimates (i.e., small confidence bounds/credible intervals), the agreement across methods improves. Despite this general agreement between methods, an important difference exists; when the rhythm amplitude is small, the state space method produces non-evolving phase estimates (e.g., near ± in Figure 4b), while phase estimates continue to evolve for the other methods.

### Fitzhugh-Nagumo oscillator

As a third example of a generative model, we consider the Fitzhugh-Nagumo oscillator, an approximation of Hodgkin-Huxley neuronal membrane potential dynamics (Izhikevich and FitzHugh, 2006). The model is defined by the following equations where (in normalized units) *V* denotes the voltage, and *I* denotes the input stimulation to the neural population (Hong 2011):

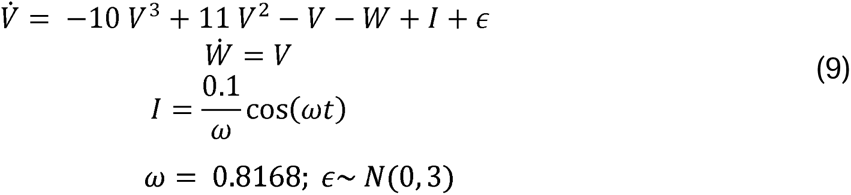

We select parameter values that permit chaotic oscillations (as elicited at a ‘strange attractor’) (Izhikevich, 2007), in which the voltage dynamics follow a trajectory consistently around a point in the coordinate space to yield a non-uniform waveform shape (relative to a sinusoid, Cole & Voytek, 2017, Figure 5a,b). This rhythm is phenomenologically similar to a slow oscillation measured in the ferret visual cortex LFP (see Figure 2A of Fröhlich & McCormick, 2010) and in hippocampal LFP in rats (see the Visual Abstract of Ouedraogo et al., 2016). After simulating data using the ode45 function in MATLAB, we rescale time such that the rhythm has an approximately 1 Hz periodicity to match a slow oscillation timescale.

**Figure 5:**
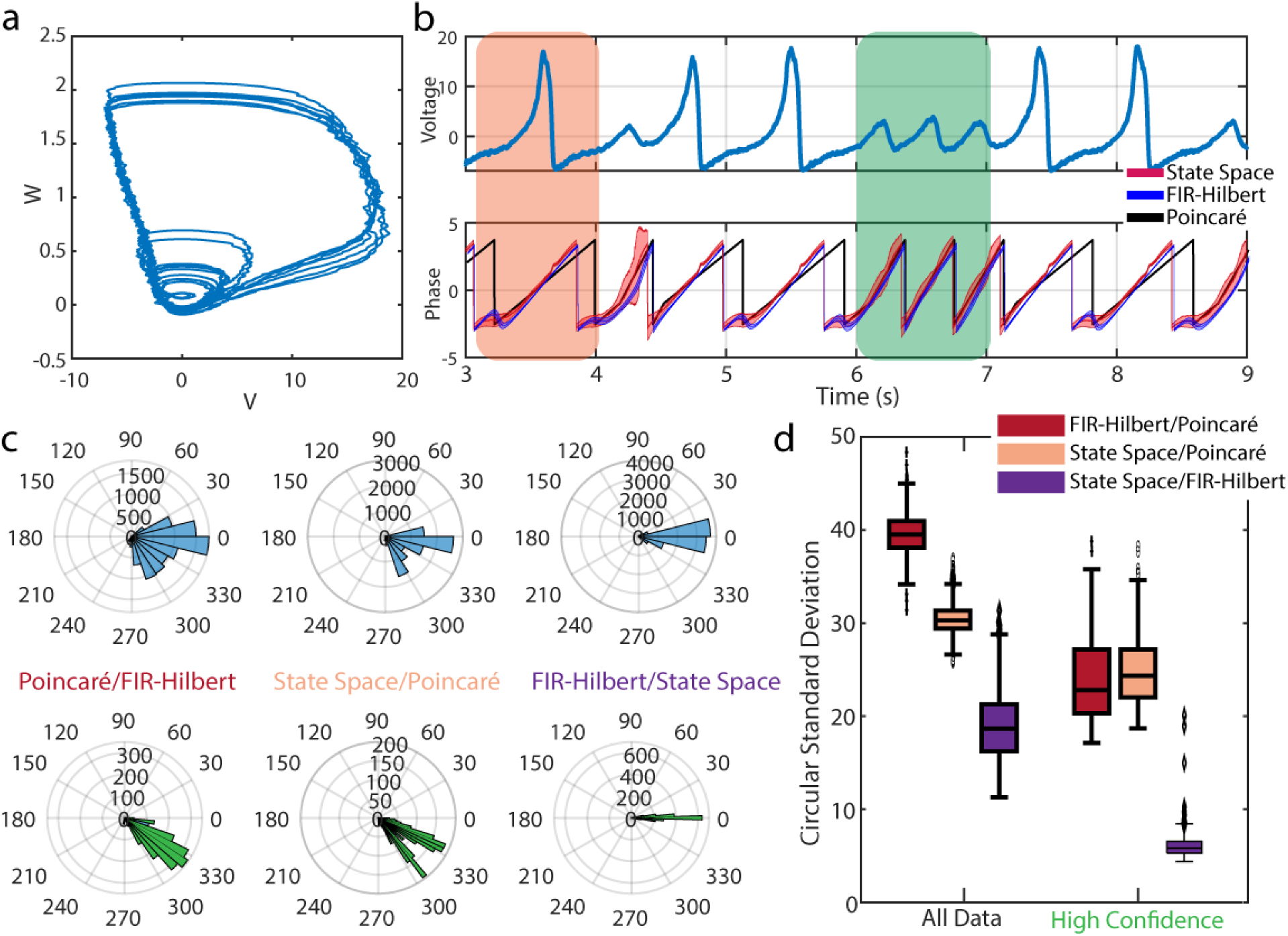
Phase estimates for a rhythm generated by the Fitzhugh-Nagumo oscillator. **(a)** Example trace of the coordinate space for the two variables comprising the Fitzhugh-Nagumo model. **(b)** (Top) Trace of example Fitzhugh-Nagumo oscillator signal. (Bottom) Phase estimates from the Poincaré method (black), FIR-Hilbert method (mean estimate blue, confidence intervals shaded blue) and state space method (mean estimate red, credible intervals shaded red). Intervals occur in which the methods agree (t *∈* [6,7]s, green rectangle) and disagree (t *∈* [3, 4] s, orange rectangle). **(c)** Differences in the estimated phase between each pair of methods. Upper row shows differences between methods across all data, and lower row shows differences only for time indices of high confidence. **(d)** Distributions of differences Distributions of differences in the estimated phase between each pair of methods across 1000 simulations using (upper) all example data or (lower) a subset of data with confident phase estimates.

In this generative model, we have access to all the variables underlying the data, i.e., the full system is known, yet we cannot define a singular phase - the definition of phase depends on the voltage events of interest. Visual inspection of the dynamics reveals waves (e.g., between 6-7 s in Figure 5b, green rectangle) for which determining whether a full wave occurs remains unclear. Two potential ways to define a waveform for this system are: (1) A repetition of the high amplitude wave (*V* > 10, see *t* near 3.5 s in Figure 3b, orange rectangle), or (2) The dynamical system completes a full cycle in the phase space at any amplitude (i.e., including the small amplitude rotations in Figure 5a, and t near 6.5 s in Figure 5b); we note that waves included in (1) are a subset of waves included in (2).

During times of smaller amplitude waves, all methods infer a complete phase cycle, and the state space and FIR-Hilbert methods suggest increased uncertainty (wider credible/confidence intervals). During higher amplitude waves, the phase cycle estimated from the Poincaré method differs from that from the other two methods (e.g., see black trace in Figure 5b in shaded orange rectangle). The Poincaré method produces a longer phase cycle for high-amplitude waves but the same phase cycle for lower amplitude waves. Additionally, we note that the state space method has wider credible intervals at the start of each phase cycle (phase at -*π*) for high-amplitude events. Comparing methods, we find more agreement between the FIR-Hilbert and state space methods (mean and SEM of the circular standard deviance across 1000 realizations of the simulation: 18.9 ± 0.1 degrees) compared to the Poincaré method (FIR-Hilbert: 39.6 ± 0.1 degrees; state space: 30.9 ± 0.05 degrees; Figure 5c,d). Differences between methods decrease when examining times with high-confidence phase estimates (FIR-Hilbert and state space: 3.04 ± 0.04; FIR-Hilbert and Poincaré: 23.9 ± 0.1; State space and Poincaré: 24.8 ± 0.1; Figure 5c,d).

We conclude that for non-sinusoidal signals with multiple waveforms, as exemplified here by the Fitzhugh-Nagumo model, phase estimates from the different methods tend to agree. As in the previous two simulation studies, considering only time-indices where we have high confidence in the phase estimates improves similarity across all methods, including to the Poincaré method. However, an important distinction between methods becomes more apparent in this simulation; the state space and FIR-Hilbert methods yield non-uniform phase cycles i.e., a non-linear ramp in the phase (see orange rectangle, Figure 5b). By contrast, the Poincaré method necessarily produces linear increases in the phase. Depending on the desired interpretation of phase – as beginning to evolve only when the voltage begins to rise (state space and FIR-Hilbert methods) as opposed to evolving purely as a function of time (Poincaré method) – different methods are appropriate.

### Confidence in phase estimates within method does not necessarily mirror phase differences across methods

In the preceding analyses, we considered both within method measures of confidence and between method differences. We now consider the importance of applying both approaches. To do so, we analyze the simulated data at different confidence/credible interval thresholds. For each threshold, we select the subset of data with confidence/credible intervals below the threshold; when the threshold is low, we are selecting only the data with the most confident phase estimates. For each threshold, we examine the cross-method phase differences for the subsets of data by computing the circular standard deviations. In some cases, we find a near linear monotonic relationship. For example, for the broadband rhythm in pink noise (Figure 6a), we find smaller deviations between methods at lower thresholds; i.e., estimate agreement across methods increases with confidence in the phase estimate. However, in other cases, we find a more complex relationship between within method confidence and between method differences. For the AR(2) and Fitzhugh-Nagumo generative models, increasing within method confidence does not necessarily decrease the between method phase difference (Figure 6b,c); in some cases, the disagreement in phase estimates between methods increases as the within method confidence increases. These results indicate that different methods can yield phase estimates that become more similar, or more dissimilar, with increasing within method confidence. This non-monotonic relationship and difference across generative models highlights the importance of considering both within method confidence/credible intervals and between method comparisons. Doing so allows a better understanding of the reliability and consistency of phase estimates across methods, as shown here for varying signal-to-noise regimes (Figure 6a,b) and nonlinear waveforms (Figure 6c).

**Figure 6.**
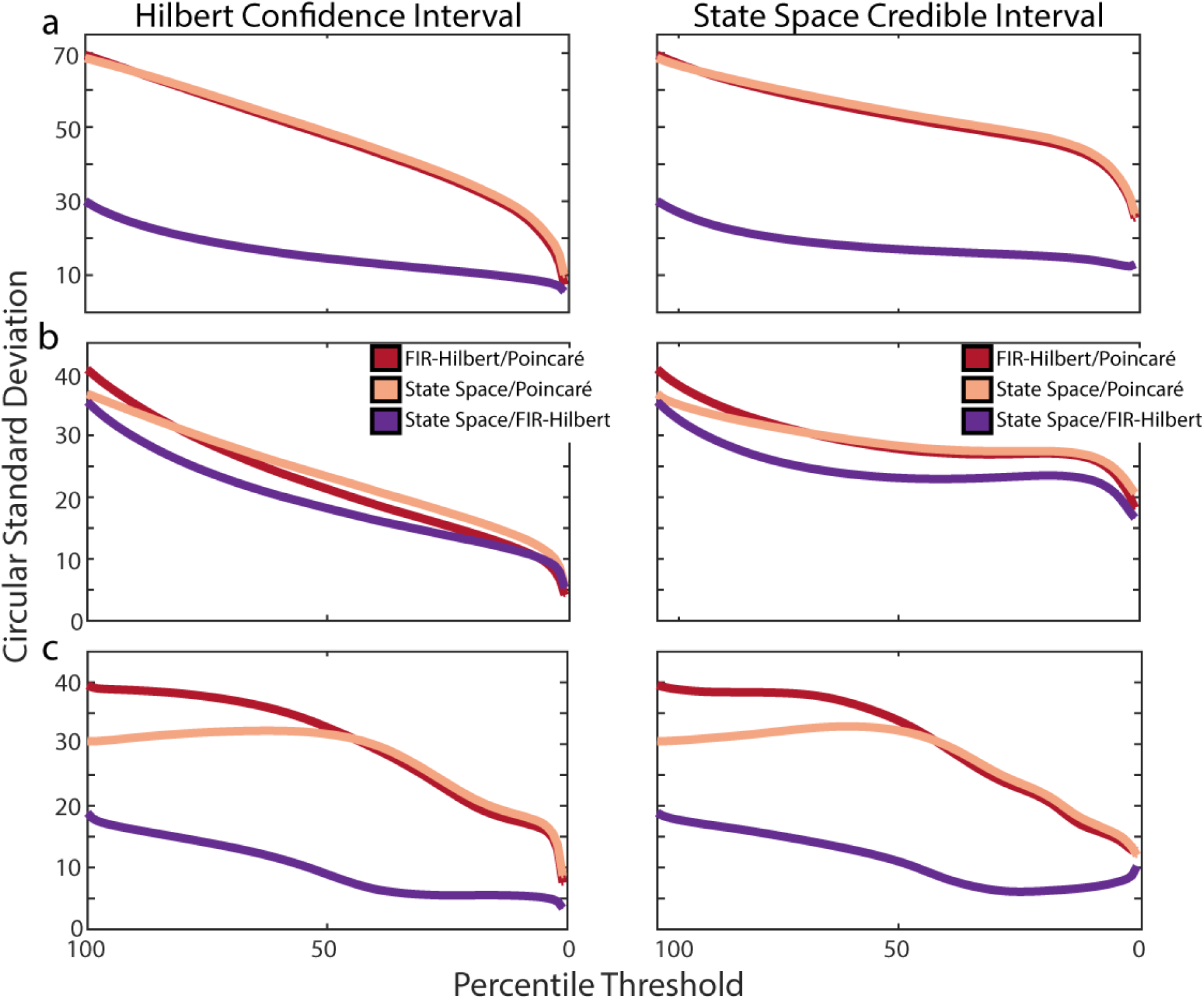
Cross-method comparisons do not directly map onto within-method confidence. **(a)** Cross-method differences for the broadband rhythm in pink noise. For each pair of methods (see legend in b) we plot the cross-method differences for different subsets of data selected based on (left column) Hilbert confidence limits or (right column) state space credible intervals. **(b)** Cross-method differences for the AR(2). **(c)** Cross-method differences for the Fitzhugh-Nagumo model.

### Example in vivo Applications

For example applications to *in vivo* data, we focus on two common analysis approaches that require phase: coherence (Pesaran et al., 2018) and phase-amplitude coupling (Hyafil et al., 2015; Tort et al., 2010) (see Methods). We analyze 400 seconds of data from two frontal electrodes with visually identifiable rhythms from an anesthetized macaque monkey (see Figure 7a, b and http://wiki.neurotycho.org/Anesthesia_Task_Details for details). Extensive analyses of these data have been conducted by Ma et al. (2019) where those authors demonstrated long range (between frontal and parietal regions) coherence in the slow delta (0.25 - 1.5 Hz) and local phase-amplitude coupling between rhythms in the slow delta (0.25 - 1.5 Hz) and beta bands (defined by Ma et al. as 15-25 Hz).

**Figure 7:**
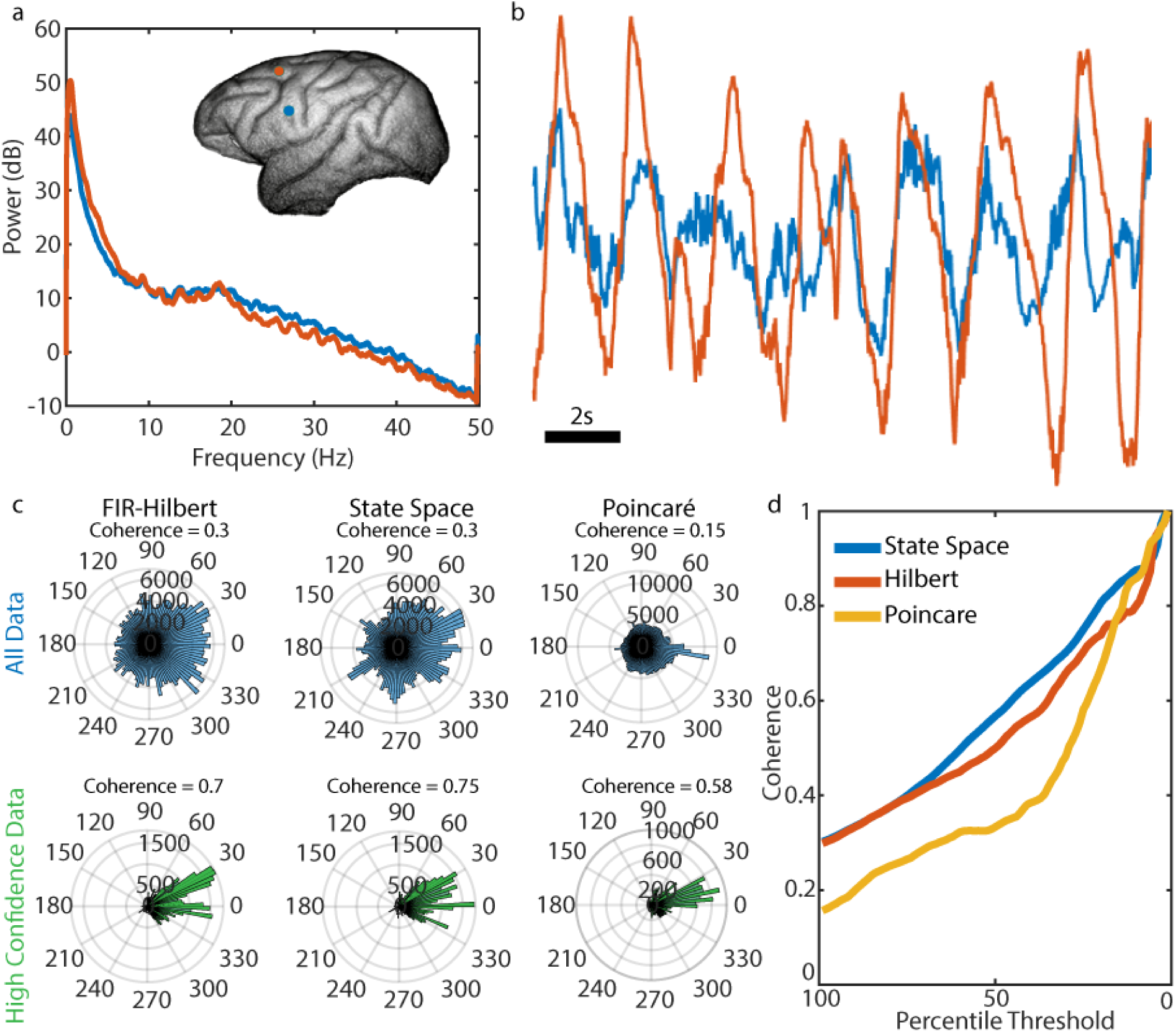
Coherence estimates increase when incorporating a measure of uncertainty in the phase. **(a)** Power spectra from multitaper estimation for the two recordings analyzed. (Inset) Macaque brain showing location of recordings. **(b)** Example traces of data from the two recordings analyzed. Note the large amplitude, slow (∼0.5 Hz) oscillation. **(c)** Estimates of coherence (numerical value) and the phase differences between recordings (histograms) under different methods and when considering all data (upper row) and only the high confidence segments of data (lower row). A bimodal distribution of phase differences appears for the FIR-Hilbert and state space methods for the high confidence segments. **(d)** Coherence estimates from subsets of data based on thresholds computed from the credible/confidence intervals.

#### Analysis of Coherence

We apply all three phase estimation methods to compute the coherence in the slow delta band. We compare coherence results from two sets of data: all the data, and a subset of the data with “high confidence” in the phase estimates. We define “high confidence” intervals as times included when thresholding for the lower quantile of confidence bounds for the FIR-Hilbert method or lower quantile of credible intervals for the state space method. Using either the FIR-Hilbert or the state space method leads to a sharp increase in the estimated coherence for the high-confidence phase data (coherence 0.7) compared to all data (coherence 0.3). We note that we cannot compare the coherence estimate using the Poincaré method to estimates from the other methods; the Poincaré method does not estimate an amplitude time series (see *Methods: Poincaré section*). However, we can compare the coherence estimated using the Poincaré method (see *Methods: Analysis of in vivo data*) across the two subsets of data. In this case, we define high confidence intervals as times at which FIR-Hilbert and state space phase estimators are simultaneously associated with low statistical uncertainty. We again find that the coherence increases from 0.15 for all data to 0.58 for the high confidence intervals of data (Figure 7c, right column). Examining the coherence estimates for subsets of data across a range of percentile thresholds for the credible/confidence intervals, we find that the coherence increases as the percentile threshold decreases; as confidence in the phase increases, so does the coherence. Because the coherence results can depend on the percentile threshold, we recommend reporting results across the range of percentile thresholds. We conclude that, for all three methods, the estimated coherence increases during time indices of high confidence phase estimates. This conclusion is not obvious; during time indices with high-confidence phase estimates, the phases and amplitudes for the slow delta rhythm could remain randomly distributed between recordings. In addition, we note that isolating the high confidence phase data produced bimodal distributions for the phase differences between electrodes with the FIR-Hilbert and state space methods, a result not clear when analyzing all data (Figure 7c). A bimodal phase-difference distribution (as seen in Figure 7c) with one mode (peak) near 0 degrees could imply a common source acting on both areas (Nunez and Srinivasan, 2006), and another mode near 30 degrees could reflect a delayed signal propagating between electrodes. We conclude that coherence estimates can be better interpreted by using segments of data with confident phase estimates.

#### Phase-Amplitude Coupling

In a second example analysis of the *in vivo* data, we use only the FIR-Hilbert and state space methods to estimate the phase-amplitude coupling between the slow delta phase and beta amplitude (see *Methods: Analysis of in vivo data*). We note that the Poincaré method is unusable for phase-amplitude coupling estimation due to the lack of an estimated amplitude time series. We expect the low-frequency delta phase to influence the high-frequency beta amplitude (Ma et al., 2019). We observe this effect when analyzing all of the data (Figure 8), and find that considering only subsets of data with high confidence in the phase estimate increases the effect size (Figure 8c) - there is greater modulation (ratio between the maximum and minimum normalized mean amplitudes across all phase bins) of the amplitude of the beta band rhythm as a function of the slow delta phase when considering only intervals with high confidence phase estimates. However, for small percentile thresholds (i.e., when confidence in the phase estimate is high, and smaller subsets of data are used for analysis), we can visually identify fluctuations in the modulation as less data is included to create the phase-amplitude plot. Like the coherence results, we recommend reporting phase-amplitude coupling metrics across the range of percentile thresholds. These results are consistent across the FIR-Hilbert and state space methods, demonstrating the robustness of this effect to model misspecification.

**Figure 8:**
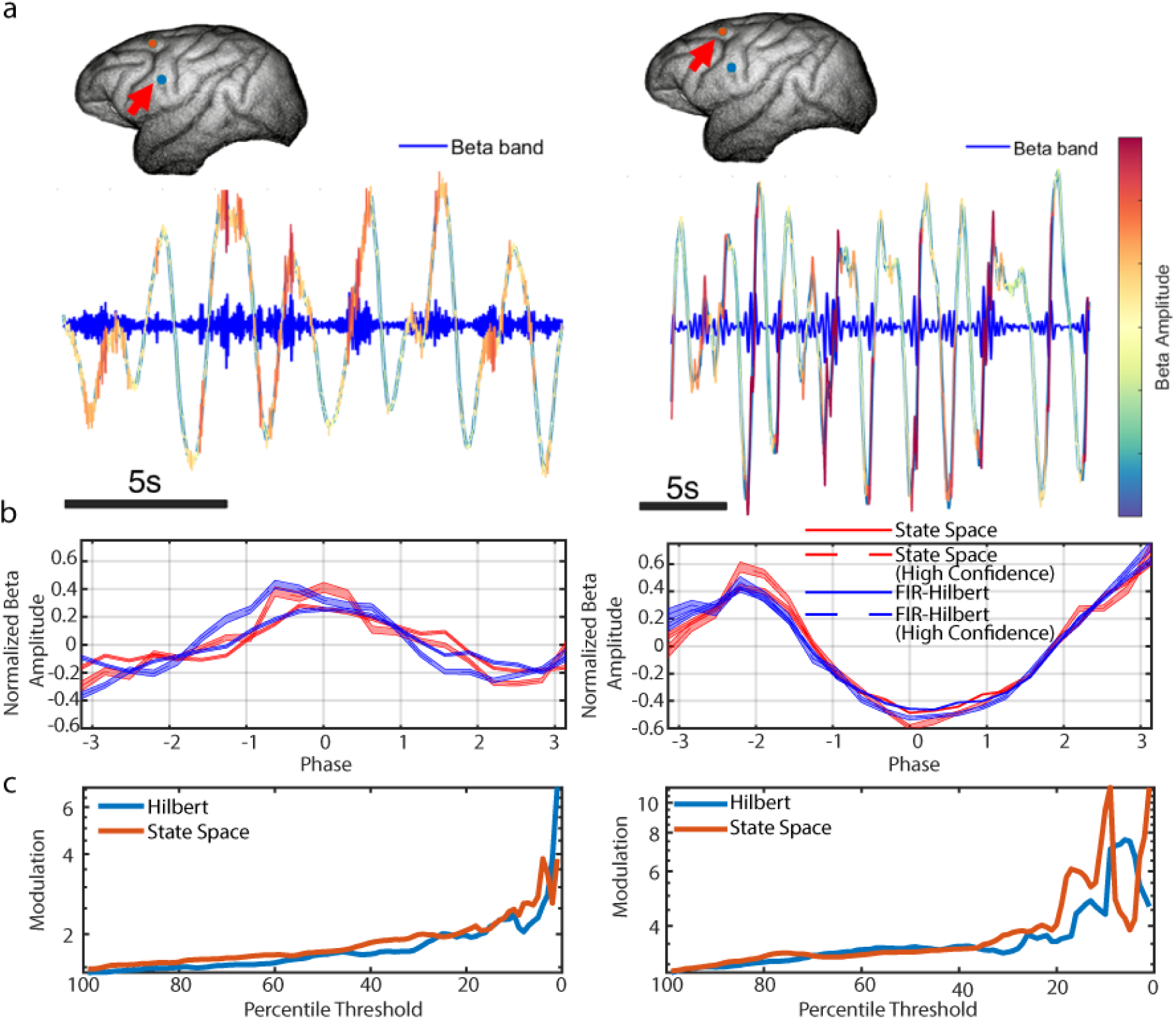
Cross-frequency coupling estimates are improved when incorporating a measure of uncertainty in the phase. **(a)** Example traces of the slow delta and beta signals showing timing of beta rhythms with specific phases of slow delta at each electrode location (inset image with arrow). Delta trace is colored according to beta amplitude (see colorbar). **(b)** Estimates of phase-amplitude coupling when using the state space method (red, shaded confidence bands) and FIR-Hilbert (blue, shaded confidence bands) method for each recording. Both methods demonstrate significant phase-amplitude coupling between slow delta phase and beta amplitude. When only using segments of data with high confidence phase (dashed curves), the modulation increases. (c) Cross-frequency coupling strength or ‘Modulation’ (the ratio between the minimum and maximum normalized mean amplitudes) estimated from subsets of data based on thresholds computed from the credible/confidence intervals.

## Discussion

In this manuscript, we demonstrated the ambiguity in specifying phase for neuroscientific applications. We used simulations from three generative models of rhythms and applied three methods to specify phase. Together, these three models and three methods encompass a wide range of empirical data and methods applied (Bruns, 2004; Cole & Voytek, 2017; Donoghue et al., 2020; Kralemann et al., 2008; Ouedraogo et al., 2016; Spyropoulos et al., 2022; Wodeyar et al., 2021). We showed that for each model of rhythms, each method produced phase estimates that optimally tracked different characteristics of the rhythms. These phase estimates converge when using only segments of data when the methods suggested greater confidence in the phase estimate. Using this insight, we applied two common analyses – coherence and phase-amplitude coupling – to example *in vivo* recordings. Considering only the subset of data with high confidence phase estimates, we found increases in both the coherence and phase-amplitude coupling, and a bimodal phase-difference distribution for the coherence that was not apparent in the original analysis. We conclude that phase is ambiguous (i.e., application specific) for neural data but using uncertainty metrics reduces the ambiguity.

Every method to specify phase imposes a model of the data time series. In practice, phase estimation is performed without careful consideration of the model (implicitly) selected. For the estimation methods considered here, we propose a decision tree to guide practitioners in choosing a phase estimator. In essence, the method chosen depends on the goal of the analysis. In the ideal case of sufficiently sampled sinusoidal data (Lainscsek et al., 2017) and no noise process, the FIR-Hilbert, state space, and Poincaré methods are all viable to identify and track inferable rhythms. If the data time series periodicity is nearly sinusoidal and noise contamination is low, then both the FIR-Hilbert and state space methods generate similar phase estimates and are not susceptible to high-frequency fluctuations like those seen using the Poincaré method. Alternatively, if there is amplitude modulation, or if the data are non-sinusoidal and noise contamination is high and the goal is to identify phase for periods with high SNR waves, then the state space method is the best option. Indeed, we note that most neural data contain noise and amplitude modulation. Finally, if the data are non-sinusoidal and the noise contamination is low, then the Poincaré method performs better at maintaining a linear ramp for the phase compared to the FIR-Hilbert and state space methods.

When phase estimation confidence is high, we typically assume that the method used to estimate phase describes the data accurately. However, high confidence does not necessarily indicate that the method accurately models the data. For instance, when a large amplitude, sudden change (i.e., a spike) occurs in the voltage time series, the Hilbert transform confidence will be high, but the FIR-Hilbert does not provide an accurate model of the data; using this method, the “phase” of the spike will depend on the filter passband. In this case, cross-method comparisons may indicate high variability across methods despite the high confidence. Low confidence in the phase estimate may occur due to an inappropriate model choice or data that do not capture identifiable rhythms (e.g., due to noise or geometry of sources). In such cases, careful inspection of the data (e.g., additional descriptive analyses to confirm the presence of a rhythm) and consideration of other phase estimation methods may help confirm if the phase can be defined as desired by the practitioner.

The features tracked by the phase are least dependent on the experimenter’s choice of estimation method when the similarity between methods is high. Across different models of rhythms, we found uncertainty metrics help in identifying segments of time when phase is similar across methods. This similarity across methods does not map linearly to the metrics of uncertainty but depends on the model studied (see *Results: Confidence in phase estimates within method does not necessarily mirror phase differences across methods*). However, the metrics of uncertainty are specific to the model assumed by each method. The uncertainty metric under the FIR-Hilbert method is a transformation of the amplitude time series for the data passed through a narrowband filter. One can identify instances when this measure of uncertainty does not represent features of the rhythm of interest. For example, when the time series is contaminated with 1/*f*^α^ noise (see *Results: Broadband Pink Noise*), high amplitude activity may appear due to noise rather than the rhythm of interest. In contrast, segments of time with low SNR rhythms can have an amplitude that fall below a given amplitude-dependent uncertainty threshold (see *Methods:* Eq. 2). Alternatively, the state space model assumes that the analytic signal representing the rhythm process is complex-Gaussian distributed and applies the Kalman filter before quantifying the uncertainty in the phase. When the signal is non-sinusoidal or the noise driving the rhythm is non-Gaussian, the credible intervals from the state space model may fail to capture confidence in the phase accurately – e.g., visible at the start of each phase cycle for high amplitude events in the Fitzhugh-Nagumo simulation (see the orange rectangle in Figure 5). Modifications of the state space model (such as the application of an extended Kalman Filter to independently track amplitude and phase or the use of particle filters - Schiff 2012) can permit more accurate modeling of non-sinusoidal rhythms and non-Gaussian noise.

While we focused on three methods to specify phase, other methods exist. We highlight a few alternatives. Sophisticated approaches to phase estimation exist in the mathematical physics literature, e.g., analysis of the mean first return time (Cao et al., 2020; Schwabedal & Pikovsky, 2013) or slowest decaying modes of the Kolmogorov backward operator (Thomas & Lindner, 2014). As these approaches are debated (Pikovsky, 2015; Thomas & Lindner, 2015), and complementary approaches developed (e.g., Engel & Kuehn, 2021), we expect their adoption and application in neuroscience will provide additional strategies to estimate phase from neural data. A recently introduced approach, the generalized phase method (Davis et al., 2022), estimates the smoothed Hilbert-transform derived phase of the largest fluctuation in the data at a given time instant. This method prioritizes the rhythm with highest amplitude present in the time series and assumes only a single rhythm is present at any instant in time. Given the similarity of this method to the basic Hilbert transform and the intent to examine distinct types of phase estimation methods, we chose not to apply this method here. Another method – the masked empirical mode decomposition or masked EMD (Quinn et al., 2021) – seeks to identify distinct waveforms in the time series. Such a method could closely track an inferable rhythm while reducing the contamination of noise and create a phase evolution that linearly tracks a non-sinusoidal rhythm. The masked EMF may struggle with identifying a singular waveform if the underlying limit-cycle attractor of the dynamical system measured is chaotic (as seen in the Fitzhugh-Nagumo simulation), leading to wave morphology that is inconsistent across cycles.

A major difficulty in identifying the best phase estimation method is that the quantity defined as phase depends on the goals of the practitioner. Thus, the idea of a “best” phase estimator is not meaningful. Instead, we recommend alternative approaches of comparing estimators, for example by comparing phase from different methods to identify consistency or testing which method of phase estimation shows the highest effect size for the feature of interest. For example, Davis and colleagues (Davis et al., 2022) compare the phase from the FIR-Hilbert method and that under the generalized phase method to the multi-unit neuronal spiking rate, demonstrating increased spike-field coherence under the generalized phase method. However, choosing the phase across methods that maximizes the relationship to neuronal activity or behavior may suffer from “voodoo correlations” (Vul et al., 2009): increases in effect size due to random chance in the method selected. Similarly, the removal of data with low confidence should be done cautiously and with a thorough understanding of the possible implications on the overall conclusions, including voodoo correlations. Further, when presenting the results, it is important to clearly communicate the limitations of the analysis and the potential impact of the removed data on the conclusions drawn. We recommend that studies present the results of phase analysis across a continuum of thresholds on the uncertainty metrics to demonstrate the implications of analyzing different subsets of data (as shown in Figure 7d and 8c). Doing so may reveal facets of the data, for example, the presence of bi-modality in the phase differences, that may otherwise be obscured by inaccurate phase estimates (see Figure 7c). The true efficacy of a method to define phase that tracks the desired feature of population neural activity (such as the peaks or troughs) could be verified by causal approaches, such as stimulation.

Closed-loop approaches to non-invasive (Klinzing et al., 2021; Ngo et al., 2013) and invasive (Fröhlich & McCormick, 2010; Lustenberger et al., 2016; Zrenner et al., 2018) stimulation based on a signal’s phase require methods for real-time phase estimation. For example, auditory stimulation triggered by the phase of a slow-wave oscillation improves memory (Ngo et al., 2013) and thalamic stimulation triggered by a patient’s tremor rhythm suppresses tremor in patients with Parkinson’s disease (Cagnan et al., 2017). The Poincaré and FIR-Hilbert methods do not directly support causal phase estimation, precluding their real-time application. However, several methods have been developed to perform real-time phase estimation using the Hilbert transform by first forecasting data under autoregressive models (Blackwood et al., 2018; Schatza et al., 2022), and a similar approach may support real-time implementation of the Poincaré method. The state space method is easily translated to a real-time framework (Wodeyar et al., 2021). Finally, similar to what we showed in the *in vivo* analyses, measures of uncertainty can help guide closed-loop stimulation to times of greater certainty in the phase estimate (Schatza et al., 2022; McNamara et al., 2022).

We note that in this manuscript we focused on the case of a single rhythm. However, the state space method supports simultaneous estimation of oscillators for multiple rhythms (Wodeyar et al., 2021; He et al., 2022). By applying the state space method to estimate multiple oscillators, we expect the state space method could provide a principled approach to phase estimation across multiple underlying oscillation patterns. The likelihood derived from the state space model can serve as an explicit indicator of which oscillation is dominant at any given time. This approach can help mitigate the risks associated with censoring data based on confidence levels and offer, for example, a more accurate representation of the coupling between spikes and neural signals in the presence of multiple rhythms. In this way, state space models may contribute to a more comprehensive analysis of the data, considering the dynamic presence and contribution of different rhythms, and ensuring that the conclusions drawn are not biased by an overemphasis on high-confidence estimates.

Phase is an important characteristic of neural activity that may be used to index cortical excitability (Freeman, 1961), availability of attentional resources (Busch et al., 2009; Fiebelkorn et al., 2019), and the neural population capacity for communication (Fries, 2015). As such, knowledge of how different phase estimators define phase is critical to address mechanistic questions in neuroscience. In this manuscript, we proposed guidelines to identify the appropriate phase estimator, namely applying estimation methods whose reconstructed phase tracks the relevant characteristics of the rhythm of interest and using metrics of statistical uncertainty. Improved approaches to phase estimation will help us better understand the network implications of rhythms and ascertain the potential for non-invasive interventions that alter cognitive ability (Ngo et al., 2013) or reduce clinical symptoms (Cagnan et al., 2017).

## Author Contributions

AW and MAK designed research, AW performed research, AW, FAM, CJC, UTE and MAK wrote the paper.

## Acknowledgements

None.

## References

1. Bernardi, G., Betta, M., Ricciardi, E., Pietrini, P., Tononi, G., & Siclari, F. (2019). Regional Delta Waves In Human Rapid Eye Movement Sleep. The Journal of Neuroscience, 39(14), 2686–2697.

2. He, B. J., Zempel, J. M., Snyder, A. Z., & Raichle, M. E. (2010). The temporal structures and functional significance of scale-free brain activity. Neuron, 66(3), 353–369.

3. Blackwood, E., Lo, M., & Widge, A. S. (2018). Continuous Phase Estimation for Phase-Locked Neural Stimulation Using an Autoregressive Model for Signal Prediction*. 2018 40th Annual International Conference of the IEEE Engineering in Medicine and Biology Society (EMBC), 4736–4739.

4. Boashash, B. (1991). Estimating and Interpreting The Instantaneous Frequency of a Signal-Part 1: Fundamentals. 19.

5. Bokil, H., Andrews, P., Kulkarni, J. E., Mehta, S., & Mitra, P. P. (2010). Chronux: A platform for analyzing neural signals. Journal of Neuroscience Methods, 192(1), 146–151.

6. Bruns, A. (2004). Fourier-, Hilbert- and wavelet-based signal analysis: Are they really different approaches? Journal of Neuroscience Methods, 137(2), 321–332.

7. Burns, S. P., Xing, D., Shelley, M. J., & Shapley, R. M. (2010). Searching for Autocoherence in the Cortical Network with a Time-Frequency Analysis of the Local Field Potential. Journal of Neuroscience, 30(11), 4033–4047.

8. Busch, N. A., Dubois, J., & VanRullen, R. (2009). The Phase of Ongoing EEG Oscillations Predicts Visual Perception. Journal of Neuroscience, 29(24), 7869–7876.

9. Cagnan, H., Pedrosa, D., Little, S., Pogosyan, A., Cheeran, B., Aziz, T., Green, A., Fitzgerald, J., Foltynie, T., Limousin, P., Zrinzo, L., Hariz, M., Friston, K. J., Denison, T., & Brown, P. (2017). Stimulating at the right time: Phase-specific deep brain stimulation. Brain, 140(1), 132–145.

10. Canolty, R. T., & Knight, R. T. (2010). The functional role of cross-frequency coupling. Trends in Cognitive Sciences, 14(11), 506–515.

11. Chavez, M., Besserve, M., Adam, C., & Martinerie, J. (2006). Towards a proper estimation of phase synchronization from time series. Journal of Neuroscience Methods, 154(1), 149–160.

12. Cohen, M. X. (2008). Assessing transient cross-frequency coupling in EEG data. Journal of Neuroscience Methods, 168(2), 494–499.

13. Cole, S. R., & Voytek, B. (2017). Brain Oscillations and the Importance of Waveform Shape. Trends in Cognitive Sciences, 21(2), 137–149.

14. Davis, Z. W., Muller, L., & Reynolds, J. H. (2022). Spontaneous Spiking Is Governed by Broadband Fluctuations. Journal of Neuroscience, 42(26), 5159–5172.

15. Donoghue, T., Haller, M., Peterson, E. J., Varma, P., Sebastian, P., Gao, R., Noto, T., Lara, A. H., Wallis, J. D., Knight, R. T., Shestyuk, A., & Voytek, B. (2020). Parameterizing neural power spectra into periodic and aperiodic components. Nature Neuroscience, 23(12), 1655–1665.

16. Fiebelkorn, I. C., Pinsk, M. A., & Kastner, S. (2019). The mediodorsal pulvinar coordinates the macaque fronto-parietal network during rhythmic spatial attention. Nature Communications, 10(1), 215.

17. Freeman, W. J. (1961). Harmonic Oscillation as Model for Cortical Excitability Changes with Attention in Cats. Science, 133(3470), 2058–2059.

18. Fries, P. (2015). Rhythms for Cognition: Communication through Coherence. Neuron, 88(1), 220–235.

19. Fröhlich, F., & McCormick, D. A. (2010). Endogenous Electric Fields May Guide Neocortical Network Activity. Neuron, 67(1), 129–143. 10.1016/j.neuron.2010.06.005

20. Herbst, S. K., Stefanics, G., & Obleser, J. (2022). Endogenous modulation of delta phase by expectation–A replication of Stefanics et al., 2010. Cortex, 149, 226–245.

21. Hong, K. S. (2011). Synchronization of coupled chaotic FitzHugh–Nagumo neurons via Lyapunov functions. Mathematics and Computers in Simulation, 82(4), 590–603.

22. Hyafil, A., Giraud, A.-L., Fontolan, L., & Gutkin, B. (2015). Neural Cross-Frequency Coupling: Connecting Architectures, Mechanisms, and Functions. Trends in Neurosciences, 38(11), 725–740.

23. Izhikevich, E. M., & Fitzhugh, R. (n.d.). FitzHugh-Nagumo model—Scholarpedia. Retrieved January 13, 2022, from http://www.scholarpedia.org/article/FitzHugh-Nagumo_model

24. Jarvis, M. R., & Mitra, P. P. (2001). Sampling properties of the spectrum and coherency of sequences of action potentials. Neural Computation, 13(4), 717–749.

25. Jones, S. R. (2016). When brain rhythms aren’t ‘rhythmic’: Implication for their mechanisms and meaning. Current Opinion in Neurobiology, 40, 72–80.

26. Klinzing, J. G., Tashiro, L., Ruf, S., Wolff, M., Born, J., & Ngo, H.-V. V. (2021). Auditory stimulation during sleep suppresses spike activity in benign epilepsy with centrotemporal spikes. Cell Reports Medicine, 2(11), 100432.

27. Kralemann, B., Cimponeriu, L., Rosenblum, M., Pikovsky, A., & Mrowka, R. (2007). Uncovering interaction of coupled oscillators from data. Physical Review E, 76(5), 055201.

28. Kramer, Mark A. “Golden Rhythms as a Theoretical Framework for Cross-Frequency Organization.” Neurons, Behavior, Data Analysis, and Theory 1 (October 20, 2022).

29. Kramer, M. A., & Eden, U. T. (2016). Case Studies in Neural Data Analysis: A Guide for the Practicing Neuroscientist. MIT Press.

30. Lachaux, J.-P., Rodriguez, E., Martinerie, J., & Varela, F. J. (1999). Measuring phase synchrony in brain signals. Human Brain Mapping, 8(4), 194–208.

31. Lepage, K. Q., Kramer, M. A., & Eden, U. T. (2011). The Dependence of Spike Field Coherence on Expected Intensity. Neural Computation, 23(9), 2209–2241.

32. Lepage, K. Q., Kramer, M. A., & Eden, U. T. (2013). Some Sampling Properties of Common Phase Estimators. Neural Computation, 25(4), 901–921.

33. Lewis, L. D., Weiner, V. S., Mukamel, E. A., Donoghue, J. A., Eskandar, E. N., Madsen, J. R., Anderson, W. S., Hochberg, L. R., Cash, S. S., Brown, E. N., & Purdon, P. L. (2012). Rapid fragmentation of neuronal networks at the onset of propofol-induced unconsciousness. Proceedings of the National Academy of Sciences, 109(49), E3377–E3386.

34. Lustenberger, C., Boyle, M. R., Alagapan, S., Mellin, J. M., Vaughn, B. V., & Fröhlich, F. (2016). Feedback-Controlled Transcranial Alternating Current Stimulation Reveals a Functional Role of Sleep Spindles in Motor Memory Consolidation. Current Biology, 26(16), 2127–2136.

35. Ma, L., Liu, W., & Hudson, A. E. (2019). Propofol Anesthesia Increases Long-range Frontoparietal Corticocortical Interaction in the Oculomotor Circuit in Macaque Monkeys. Anesthesiology, 130(4), 560–571.

36. Matsuda, T., & Komaki, F. (2017). Time Series Decomposition into Oscillation Components and Phase Estimation. Neural Computation, 29(2), 332–367.

37. McNamara, C. G., Rothwell, M., & Sharott, A. (2022). Stable, interactive modulation of neuronal oscillations produced through brain-machine equilibrium. Cell Reports, 41(6), 111616.

38. Miller, K. J., Zanos, S., Fetz, E. E., Den Nijs, M., & Ojemann, J. G. (2009a). Decoupling the cortical power spectrum reveals real-time representation of individual finger movements in humans. Journal of Neuroscience, 29(10), 3132–3137.

39. Miller, K. J., Sorensen, L. B., Ojemann, J. G., & Den Nijs, M. (2009b). Power-law scaling in the brain surface electric potential. PLoS computational biology, 5(12), e1000609.

40. Morey, R. D., Hoekstra, R., Rouder, J. N., Lee, M. D., & Wagenmakers, E.-J. (2016). The fallacy of placing confidence in confidence intervals. Psychonomic Bulletin & Review, 23(1), 103–123.

41. Mormann, F., Lehnertz, K., David, P., & Elger, C. E. (2000). Mean phase coherence as a measure for phase synchronization and its application to the EEG of epilepsy patients. Physica D: Nonlinear Phenomena, 144(3-4), 358–369.

42. Nadalin, J. K., Martinet, L. E., Blackwood, E. B., Lo, M. C., Widge, A., Cash S. S., Eden U. T., & Kramer, M. A. (2019). A statistical framework to assess cross-frequency coupling while accounting for confounding analysis effects. Elife, 8, e44287.

43. Navas-Olive, A., Valero, M., Jurado-Parras, T., de Salas-Quiroga, A., Averkin, R. G., Gambino, G., Cid, E., & de la Prida, L. M. (2020). Multimodal determinants of phase-locked dynamics across deep-superficial hippocampal sublayers during theta oscillations. Nature Communications, 11(1), Article 1.

44. Ngo, H.-V. V., Martinetz, T., Born, J., & Mölle, M. (2013). Auditory Closed-Loop Stimulation of the Sleep Slow Oscillation Enhances Memory. Neuron, 78(3), 545–553.

45. Nunez, P. L., & Srinivasan, R. (2006). Electric fields of the brain: the neurophysics of EEG. Oxford University Press, USA.

46. Ouedraogo, D. W., Lenck-Santini, P.-P., Marti, G., Robbe, D., Crépel, V., & Epsztein, J. (2016). Abnormal UP/DOWN Membrane Potential Dynamics Coupled with the Neocortical Slow Oscillation in Dentate Granule Cells during the Latent Phase of Temporal Lobe Epilepsy. Eneuro, 3(3), ENEURO.0017-16.2016.

47. Pesaran, B., Vinck, M., Einevoll, G. T., Sirota, A., Fries, P., Siegel, M., Truccolo, W., Schroeder, C. E., & Srinivasan, R. (2018). Investigating large-scale brain dynamics using field potential recordings: Analysis and interpretation. Nature Neuroscience, 21(7), Article 7.

48. Petersen, P. C., & Buzsáki, G. (2020). Cooling of Medial Septum Reveals Theta Phase Lag Coordination of Hippocampal Cell Assemblies. Neuron, 107(4), 731–744.e3.

49. Quinn, A. J., Lopes-dos-Santos, V., Huang, N., Liang, W.-K., Juan, C.-H., Yeh, J.-R., Nobre, A. C., Dupret, D., & Woolrich, M. W. (2021). Within-cycle instantaneous frequency profiles report oscillatory waveform dynamics. Journal of Neurophysiology, 126(4), 1190–1208.

50. Ray, S., Crone, N. E., Niebur, E., Franaszczuk, P. J., & Hsiao, S. S. (2008). Neural Correlates of High-Gamma Oscillations (60–200 Hz) in Macaque Local Field Potentials and Their Potential Implications in Electrocorticography. The Journal of Neuroscience, 28(45), 11526–11536.

51. Roberts, M. J., Lowet, E., Brunet, N. M., Ter Wal, M., Tiesinga, P., Fries, P., & De Weerd, P. (2013). Robust Gamma Coherence between Macaque V1 and V2 by Dynamic Frequency Matching. Neuron, 78(3), 523–536.

52. Rosenblum, M. G., Pikovsky, A. S., & Kurths, J. (1997). Phase synchronization in driven and coupled chaotic oscillators. IEEE Transactions on Circuits and Systems I: Fundamental Theory and Applications, 44(10), 874–881.

53. Schatza, M. J., Blackwood, E. B., Nagrale, S. S., & Widge, A. S. (2022). Toolkit for Oscillatory Real-time Tracking and Estimation (TORTE). Journal of Neuroscience Methods, 366, 109409.

54. Schiff, S. J. (2012). Neural control engineering: The emerging intersection between control theory and neuroscience. MIT Press.

55. Shumway, R. H., & Stoffer, D. S. (1982). An Approach to Time Series Smoothing and Forecasting Using the Em Algorithm. Journal of Time Series Analysis, 3(4), 253–264.

56. Shumway, R. H., & Stoffer, D. S. (2017). ARIMA Models. In R. H. Shumway & D. S. Stoffer (Eds.), Time Series Analysis and Its Applications: With R Examples (pp. 75–163). Springer International Publishing.

57. Soulat, H., Stephen, E. P., Beck, A. M., & Purdon, P. L. (2022). State space methods for phase amplitude coupling analysis. Scientific Reports, 12(1), Article 1.

58. Spyropoulos, G., Saponati, M., Dowdall, J. R., Schölvinck, M. L., Bosman, C. A., Lima, B., Peter, A., Onorato, I., Klon-Lipok, J., Roese, R., Neuenschwander, S., Fries, P., & Vinck, M. (2022). Spontaneous variability in gamma dynamics described by a damped harmonic oscillator driven by noise. Nature Communications, 13(1), Article 1.

59. Takens, F. (1981). Detecting strange attractors in turbulence. In D. Rand & L.-S. Young (Eds.), Dynamical Systems and Turbulence, Warwick 1980 (pp. 366–381). Springer.

60. Tort, A. B. L., Komorowski, R., Eichenbaum, H., & Kopell, N. (2010). Measuring Phase-Amplitude Coupling Between Neuronal Oscillations of Different Frequencies. J Neurophysiol, 104, 16.

61. VanRullen, R. (2016). Perceptual Cycles. Trends in Cognitive Sciences, 20(10), 723–735.

62. Vul, E., Harris, C., Winkielman, P., & Pashler, H. (2009). Puzzlingly High Correlations in fMRI Studies of Emotion, Personality, and Social Cognition. Perspectives on Psychological Science, 4(3), 274–290.

63. Widge, A. S., & Miller, E. K. (2019). Targeting Cognition and Networks Through Neural Oscillations: Next-Generation Clinical Brain Stimulation. JAMA Psychiatry, 76(7), 671–672.

64. Widmann, A., Schröger, E., & Maess, B. (2015). Digital filter design for electrophysiological data—A practical approach. Journal of Neuroscience Methods, 250, 34–46.

65. Wodeyar, A., Schatza, M., Widge, A. S., Eden, U. T., & Kramer, M. A. (2021). A state space modeling approach to real-time phase estimation. ELife, 10, e68803.

66. Wodeyar, A., & Srinivasan, R. (2022). Structural connectome constrained graphical lasso for MEG partial coherence. Network Neuroscience, 6(4), 1219–1242.

67. Xing, D., Shen, Y., Burns, S., Yeh, C.-I., Shapley, R., & Li, W. (2012). Stochastic Generation of Gamma-Band Activity in Primary Visual Cortex of Awake and Anesthetized Monkeys. Journal of Neuroscience, 32(40), 13873–13880a.

68. Yael, D., Vecht, J. J., & Bar-Gad, I. (2018). Filter-Based Phase Shifts Distort Neuronal Timing Information. ENeuro, 5(2).

69. Zanos, S., Zanos, T. P., Marmarelis, V. Z., Ojemann, G. A., & Fetz, E. E. (2012). Relationships between spike-free local field potentials and spike timing in human temporal cortex. Journal of Neurophysiology, 107(7), 1808–1821.

70. Zhong, W., Ciatipis, M., Wolfenstetter, T., Jessberger, J., Müller, C., Ponsel, S., Yanovsky, Y., Brankačk, J., Tort, A. B. L., & Draguhn, A. (2017). Selective entrainment of gamma subbands by different slow network oscillations. Proceedings of the National Academy of Sciences, 114(17), 4519–4524.

71. Zrenner, C., Desideri, D., Belardinelli, P., & Ziemann, U. (2018). Real-time EEG-defined excitability states determine efficacy of TMS-induced plasticity in human motor cortex. Brain Stimulation, 11(2), 374–389.

72. Zrenner, C., Dragana Galevska, Jaakko O. Nieminen, David Baur, MariaIoanna Stefanou, & Ulf Ziemann. (2020). The shaky ground truth of real-time phase estimation. 21.

